# Semi-processive hyperglycosylation of adhesin by bacterial protein *N*-glycosyltransferases

**DOI:** 10.1101/2020.09.10.281741

**Authors:** Liubov Yakovlieva, Carlos Ramírez-Palacios, Siewert J. Marrink, Marthe T.C. Walvoort

## Abstract

Processivity is an important feature of enzyme families such as DNA polymerases, polysaccharide synthases and protein kinases, to ensure high fidelity in biopolymer synthesis and modification. Here we reveal processive character in the family of cytoplasmic protein *N*-glycosyltransferases (NGTs). Through various activity assays, intact protein mass spectrometry and proteomics analysis, we established that NGTs from non-typeable *Haemophilus influenzae* and *Actinobacillus pleuropneumoniae* modify an adhesin protein fragment in a semi-processive manner. Molecular modeling studies suggest that the processivity arises from the shallow substrate binding groove in NGT, that promotes the sliding of the adhesin over the surface to allow further glycosylations without temporary dissociation. We hypothesize that the processive character of these bacterial protein glycosyltransferases is the mechanism to ensure multisite glycosylation of adhesins *in vivo*, thereby creating the densely glycosylated proteins necessary for bacterial self-aggregation and adherence to human cells, as a first step towards infection.

## Introduction

Protein glycosylation is a ubiquitous post-translation modification wherein amino acid side chains of proteins are decorated with carbohydrates. Glycosylation affects many properties of the modified protein (*e*.*g*. solubility, stability, transport) and influences the biochemical pathways the glycoprotein is involved in, such as signalling, communication, and interaction with receptors.^1^ Interestingly, the majority of glycoproteins feature complex glycans attached at specific positions (*e*.*g*. antibodies), and their truncation or absence can greatly influence the function of the glycoprotein and the downstream processes (*e*.*g*. in cancer).^2^ On the other hand, there are examples of glycoproteins where the sheer number of carbohydrate modifications seems to be more important for biological activity than the specific location. For instance, in the case of mucins, several *O*-GalNAc-transferases, each with specific substrate specificity, work in concert to create a densely covered glycan surface.^3^ In bacteria, an increasing number of proteins are known to be densely glycosylated (hyperglycosylated), and these proteins are often involved in virulence traits such as adhesion and autoaggregation.^4^

Little is known about the mechanistic aspects of protein hyperglycosylation (or multisite glycosylation), and how protein glycosyltransferases (GTs) control the efficiency of surface modification. The majority of the biosynthetic processes that produce glycoproteins can broadly be divided into two categories, *i*.*e*. enzymes involved in *N*-glycosylation that transfer a pre-assembled lipid-linked glycan *en bloc* to an asparagine residue in the consensus sequence N*-X*-(S/T) (where *X* ≠ Pro), such as the well-known eukaryotic OST complex^5^ and its bacterial homologue PglB,^6^ and enzymes responsible for *O*-linked glycosylation, that transfer single carbohydrate residues from soluble nucleotide-activated substrates to serine and threonine, such as *O*-GlcNAc transferase (OGT)^7^ and *O*-GalNAc transferases involved in the initiation of mucin glycosylation.^3^ *N*-linked glycosylation occurs predominantly co-translationally on a limited number of residues, and subsequent trimming and/or further modification of the glycan results in a tremendous diversity in glycoforms, as exemplified by the >200 erythropoietin glycoforms identified in a single sample.^8^ On the other hand, *O*-linked glycosylation mostly happens post-translationally, and is often driven by nucleotide-sugar substrate concentrations.^9^

An intriguing glycosylation system that combines characteristics of both categories is the family of cytoplasmic *N*-glycosyltransferases (NGT), which is unique to bacteria. The first NGT, called HMW1C, was identified in non-typeable *Haemophilus influenzae* (NTHi),^10,11^ and is responsible for the multisite glycosylation of high-molecular weight (HMW) adhesin HMW1A. Together with the translocator HMW1B, this two-partner secretion system produces densely glycosylated adhesins on the extracellular surface of NTHi, which are crucial for adherence to human epithelial cells, as the first step in infection. Soon after this first report, homologous NGTs were identified in *Actinobacillus pleuropneumoniae*,^12^ *Yersinia enterocolitica*,^13^ *Kingella kingae*, and *Aggregatibacter aphrophilus*.^14^ NGTs generally catalyze the transfer of a single glucose (Glc) residue from the nucleotide-activated donor UDP-α-D-Glc to an asparagine residue in the consensus sequence (N-X-S/T). They are metal-independent inverting GTs, creating a β-linked modification, and based on structural similarities are classified in GT family 41 (CAZy database),^15,16^ together with the soluble *O*-GlcNAc transferase (OGT) as the only other member. Interestingly, NGTs display a relaxed sequence requirement, as modification on non-sequon Asn residues, and modification on residues other than Asn have been observed.^17^ Moreover, also di-hexose modifications have been identified both *in vivo* and *in vitro*, suggesting that NGTs may have the ability to generate both protein *N*-linkages, and glycan *O*-linkages.^10,18^ The majority of known acceptor substrates of NGTs belong to the class of adhesins and autotransporters, which are generally large membrane-associated proteins that play a distinct role in virulence.^19,20^ It is noteworthy that in almost all examples where *N*-linked glucosylation activity was confirmed, a large number of glucose moieties was added to the native protein substrates.^17,18^ The importance of multisite glycosylation for adherence was confirmed when heterologous co-expression of KkNGT and Knh in a non-adherent *E. coli* resulted in bacterial adherence to human epithelial cells.^14^

To unravel the mechanism of bacterial multisite protein glycosylation, we questioned whether hyperglycosylation is the result of a processive mechanism in NGT. This research question was inspired by the fast modification by ApNGT of the *C*-terminal fragment of HMW1A adhesin that we observed when producing *in vitro* glucosylated adhesin fragments for antibody binding studies.^21^ Processivity is a complex mechanistic feature that has been identified in a variety of enzymes, including DNA polymerases, ubiquitin ligases, protein kinases, and enzymes involved in polysaccharide synthesis and breakdown (glycosyl transferases and hydrolases),^22^ but has not yet been identified in protein GTs. In a processive mechanism, NGT would modify the adhesin substrate with multiple glucoses during a single substrate binding event (**Figure 1A**). Because multiple rounds of catalysis happen before dissociation, a processive mechanism would result in the fast generation of multiply glycosylated proteins. Alternatively, NGT may employ a distributive mechanism, in which every binding event is followed by glucose transfer and release of the resulting product (**Figure 1B**). For a subsequent modification, the adhesin substrate has to bind again, and as a result, modifications would be introduced in a stepwise manner and products reflect a distribution of modifications. A distributive mechanism has been observed for the OGT-catalyzed O-GlcNAcylation of RNA polymerase II.^23^ Processivity is a challenging trait to study, and established methods have been reviewed elsewhere.^22,24^We selected HiNGT (R2846_0712) and its close homolog ApNGT (APL_1635, 65% identity and 85% similarity)^12^ and using the C-terminal region of the natural HMW1A adhesin (HMW1ct, from *H. influenzae*, **Figure S1**) as acceptor substrate, we show that both NGTs display semi-processive behaviour (**Figure 1C**). Moreover, using molecular dynamics simulations we provide insight into the structural factors that may be at the basis of adhesin hyperglycosylation. Our research establishes a novel mechanism in the family of protein *N*-glycosyltransferases, that will advance our understanding of bacterial hyperglycosylation and is important for the application of the NGT system in glycoprotein production.

**Figure 1.**
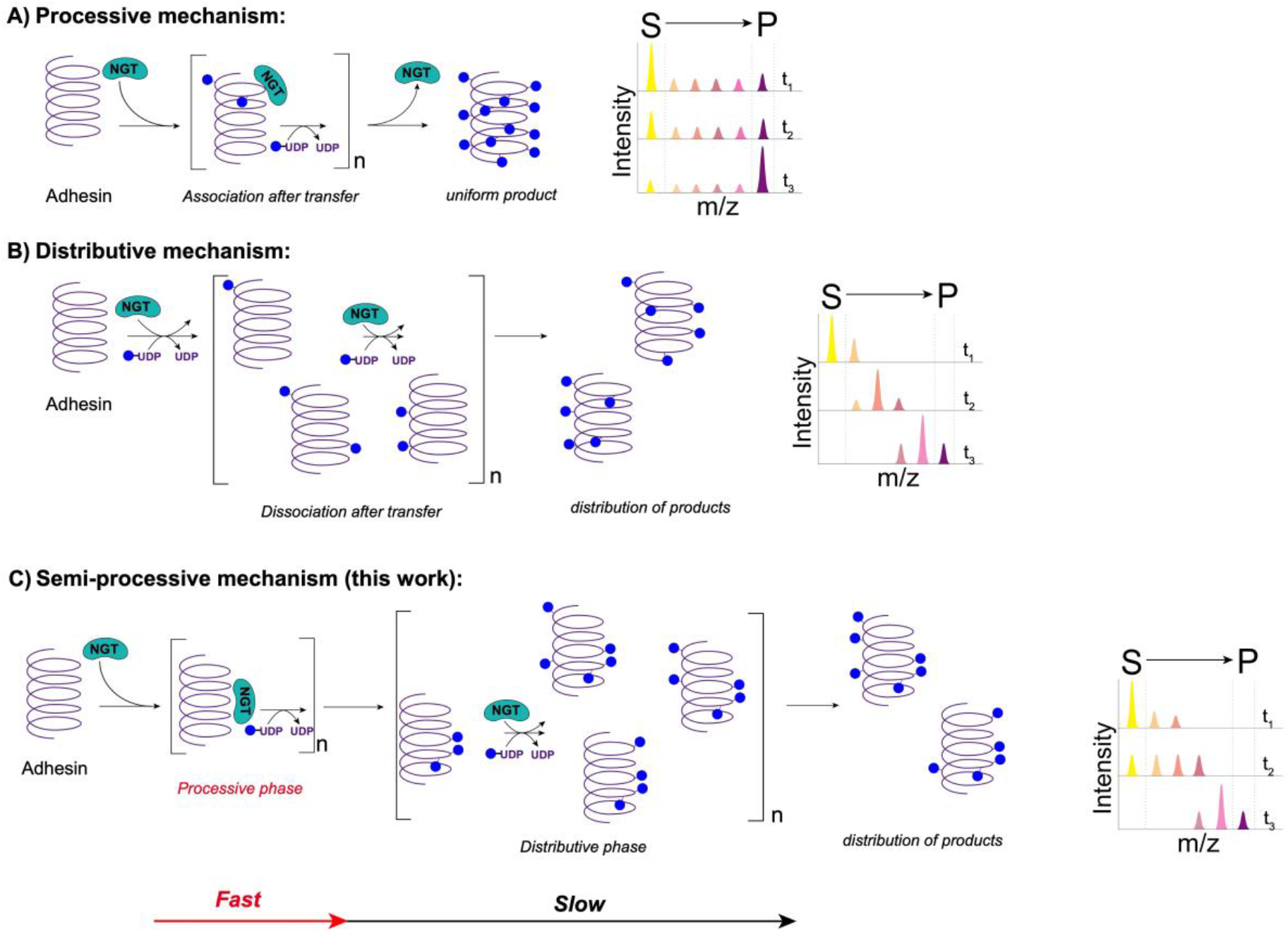
Schematic representation of the mechanism and product profiles in (**A**) a processive mechanism, (**B**) a distributive mechanism, and (**C**) the semi-processive mechanism of adhesin hyperglycosylation proposed in this work. NGT = *N*-glycosyltransferase, dark blue circles = glucose.

## Results

### Glycosylation of HMW1ct proceeds via an initial fast processive phase

To get a first impression of the glycosylation efficiency on the adhesin substrate HMW1ct, the reaction by ApNGT and HiNGT was monitored over time by examining the product profiles. *In vitro* reactions were performed at room temperature with varying enzyme to substrate ratios and quenched at certain time points by heating to 100 °C for 10 minutes. As depicted in **Figure 2A** when the ratio ApNGT to HMW1ct adhesin was 1:10, glycosylation occurred rapidly and led to the formation of a mixture of 3-6 times glucosylated (3-Glc to 6-Glc) product within 5 minutes. Over the next 15 hours, this 6-Glc product was slowly but steadily converted to even higher-order glycoforms (7-Glc and 8-Glc). Interestingly, in the first minute of the reaction no significant accumulation of a single early glycoform was observed, but rather a broad distribution of 1-Glc to 4-Glc products. Moreover, low levels of the substrate and early glycoforms (0-Glc to 2-Glc) persisted in the first 10 minutes. To slow down the rate of product formation and capitalize on intrinsic binding affinity instead of concentration effects, the experiment was repeated with a ratio of ApNGT to HMW1ct adhesin of 1:100 (**Figure 2B**). The product profile thus obtained provided a more pronounced effect, in which early and intermediate glycoforms are rapidly produced, resulting in low level accumulation of intermediate products (1-Glc to 6-Glc) in 10 minutes (**Figure 2C**), which are subsequently converted to 7-Glc and 8-Glc as the major products after 15 hours. The absence of significant levels of one intermediate glycoform before 30 minutes is intriguing, as is the persistence of non-modified substrate (0-Glc) while advanced glycoforms are being produced. While the adhesin substrate is present in large access (enzyme:substrate is 1:100), especially at the beginning of the reaction, it appears that for ApNGT formation of the first glycoform triggers the production of the next one in a processive manner. Using a continuous assay that quantifies UDP release, a clear transition from the fast phase to the slow phase was also observed (**Figure 2G**). Close inspection of the progress curve of ApNGT:HMW1ct 1:100 reveals a short ‘lag-phase’ in the first minutes, where the rate of UDP formation quickly increases, indicative of the increasing affinity of ApNGT for the early glycoform products. In an attempt to quantify this early processive behaviour, the processivity factor *P*_*n*_ was calculated using the profile at 10 minutes (**Figure 2I, Table S1**). The *P*_*n*_ value reflects the probability that the enzyme will remain associated to add an additional modification (n+1) instead of dissociating.^25,26^ The *P*_*n*_ value for the first addition was 0.22, which suggests that only 22% of ApNGT that added the first glucose continued on to add more modifications. Intriguingly, the *P*_*n*_ values for the next two additions were high (0.92 and 0.95, respectively), revealing that the production of the 3-Glc and 4-Glc products happens with considerable processivity. Subsequently, the *P*_*n*_ value drops to 0.74 (for 5-Glc) and 0.34 (for 6-Glc), which supports a change to a more distributive mechanism.

**Figure 2.**
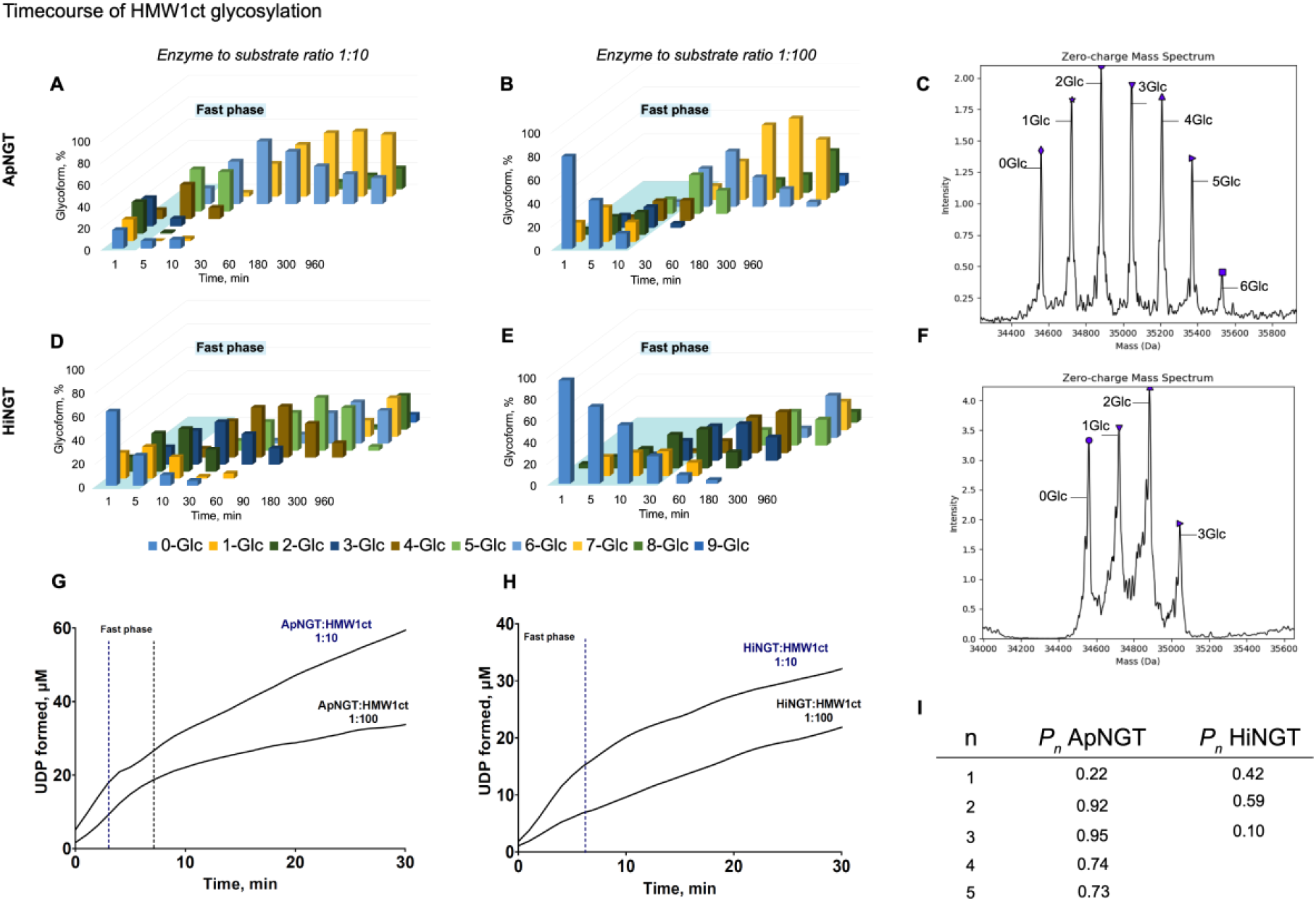
Time-course experiments and kinetic parameters of the glycosylation reaction of HMW1ct with ApNGT and HiNGT. A) Time-course product profile of ApNGT and HMW1ct in a ratio of 1:10; B) Time-course product profile of ApNGT and HMW1ct in a ratio of 1:100; C) Deconvolved mass spectrum of the product profile generated from ApNGT:HMW1ct 1:100 at 10 minutes; D) Time-course product profile of HiNGT and HMW1ct in a ratio of 1:10; E) Time-course product profile of HiNGT and HMW1ct in a ratio of 1:100; F) Deconvolved mass spectrum of the product profile generated from HiNGT:HMW1ct 1:10 at 5 minutes; For panels A-F, every reaction contained 10µM of HMW1ct protein substrate and the molarity of the enzyme was adjusted according to the desired ratio. UDP-Glc is present in excess (1mM). Representative data of two independent experiments are shown. Deconvolved spectra for selected time-points are available in Supplementary **Figures S2-S5**. The light blue panel highlights the processive fast phase; G) Reaction progress continuously monitored with the coupled-assay for ApNGT; H) Reaction progress continuously monitored with the coupled-assay for HiNGT; I) Processivity parameters obtained for ApNGT and HiNGT.

The HiNGT-catalyzed HMW1ct glycosylation appears to produce product profiles that share characteristics with the profiles from ApNGT, however the trend is less pronounced and develops at a significantly slower rate. When the reaction was performed with a ratio of HiNGT to adhesin of 1:10 (**Figure 2D**), a broad distribution of glycoforms (1-Glc to 3-Glc) was formed in the first 5 minutes (**Figure 2F**). Subsequently, these glycoforms were gradually further modified to reach mixtures where the major products were 2-Glc and 3-Glc (10 minutes), 3-Glc and 4-Glc (30 minutes), 4-Glc and 5-Glc (90 minutes), and 5-Glc and 6-Glc (300 minutes). After 15 hours the final glycoforms contained mostly 7-9 Glc moieties. This period in which a batch of glycoforms is collectively modified to produce more substituted products yields a product profile that resembles a Poisson distribution,^27^ which is associated with a distributive mechanism. Performing the reaction with a ratio of HiNGT to adhesin of 1:100 (**Figure 2E**) again emphasized the processive behaviour in the first phase, where early glycoforms are rapidly generated while non-modified substrate (0-Glc) persists for at least 180 minutes. Progress curves obtained with the continuous coupled-assay again indicate a change from a fast phase to a slow phase, especially for a ratio of HiNGT:HMW1ct 1:10 (**Figure 2H**). In the case of HiNGT, the *P*_*n*_ parameters (at 30 minutes, **Figure 2I, Table S2**) for the first additions were 0.42 (to 2-Glc), 0.59 (to 3-Glc) and 0.10 (to 4-Glc), suggesting that most processive character was displayed at the addition of the third glucose.

To quantify the difference in reaction kinetics between ApNGT and the slower HiNGT, we determined *k*_cat_ and *K*_m_ using the continuous coupled-assay (**Figure S6**). ApNGT followed typical Michaelis-Menten kinetics, which has been linked to processive character in the case of multisite phosphorylation, resulting in *k*_cat_ 0.74 −0.99 s^-1^ and *K*_m_ 6.09 – 15.6 μM.^28,29^ In contrast, for HiNGT the initial velocities (*V*_0_) were found to increase linearly and did not reach a maximum level at the highest HMW1ct concentration (**Figure S7**). This suggests that the activity of HiNGT is more dependent on the HMW1ct concentration than is the case for ApNGT. In addition, we postulate that especially in the case of HiNGT, higher HMW1ct concentrations lead to a fast production of inhibitory products (*vide infra*). In analogy to studies on multisite phosphorylation,^30^ this product inhibition may stem from a more distributive character. These experiments together paint a picture in which ApNGT, in particular, displays processive behaviour in the initial fast phase, followed by a transition to a slower phase with more distributive characteristics. HiNGT seems to follow the same trend, albeit with a shorter fast processive phase.

### Product inhibition causes a mechanism change and determines the final product profile

With the production of 5-Glc and 6-Glc for ApNGT (30 min, **Figure 2B**) and 2-Glc and 3-Glc for HiNGT (60 min, **Figure 2E**), the reaction seems to enter into a slow phase that has a more distributive character. Because it was observed previously that ApNGT has a high affinity for the Glc-adhesin product, which seriously hampered the purification by standard methods,^17,21^ we hypothesized that this mechanistic transition was due to a competing binding of the glycosylated products. The affinity of ApNGT towards substrate (HMW1ct) or product (Glc-HMW1ct, mixture of 7,8,9,10-Glc glycoforms) was determined using surface plasmon resonance (SPR). Interestingly, the *K*_D_ values were in the same range (HMW1ct *K*_D_ 5.85 ± 4.49 μM, Glc-HMW1ct *K*_D_ 9.81 ± 1.55 μM), suggesting that ApNGT binds both the substrate and the product with equal affinity (**Figure S8**). Unfortunately, we were not able to perform the same studies with HiNGT, as concentrated solutions of the enzyme were not stable enough for SPR experiments.

Based on the similar affinities of ApNGT for both the adhesin substrate (HMW1ct) and product (Glc-HMW1ct), we set out to evaluate the influence of concentration on the extent of glycosylation. We hypothesized that if the production of glycosylated product interferes with the efficiency of the reaction, increasing the substrate concentration will enhance the production of these inhibitory glycoforms, resulting in an overall reduced glycosylation efficiency. This effect has been observed before in an *ex vivo* expression system of HiNGT and HMW1A (full-length *H. influenzae* adhesin), where the increasing expression of HMW1A resulted in a reduction of site-specific glycan occupancy.^31^ We performed overnight glycosylation reactions in which the ratio ApNGT:HMW1ct was kept constant at 1:100, and the ratio of HiNGT:HMW1ct at 1:10, while the concentration of HMW1ct was varied from 5 μM to 100 μM (**Figure S9**), and the UDP-Glc concentration was fixed at 1 mM. Indeed, upon increasing the concentration of HMW1ct in the ApNGT-catalyzed reaction, the final distribution of glycoforms reduced from 7-Glc to 9-Glc (5 μM HMW1ct) to 1-Glc to 5-Glc (100 μM HMW1ct). A similar trend was observed for HiNGT, although the efficiency at the lowest HMW1ct concentration (5 μM) was also greatly reduced, presumably because of the fine balance between glycosylation and inhibition of the catalytically poor HiNGT at low concentrations. Interestingly, the inhibitory effect was greatly diminished when the concentration of UDP-Glc was increased proportionally to HMW1ct (**Figure S10**). For both ApNGT and HiNGT, product profiles (6-Glc and 7-Glc for ApNGT, 7-Glc to 9-Glc for HiNGT) close to the fully glycosylated distribution were again observed.

### Glycosylated HMW1ct inhibits processivity, while early glycoforms efficiently alleviate inhibition

To obtain a better understanding of the processive fast phase of HMW1ct glycosylation, and the influence of glycosylated adhesin on processivity, a distraction assay was performed. The principle of this experiment is to test the ability of a competitor, which is typically an inhibitor or a new batch of (labelled) substrate, to distract the processive enzyme from the substrate it is associated with. Since there are no known inhibitors of NGT glycosyltransferases, we decided to make use of the high affinity of the NGT enzymes for their glycosylated products (*vide supra*), called Glc-HMW1ct (mixture of 7,8,9,10-Glc glycoforms). Intriguingly, when the ApNGT-HMW1ct reaction (ratio 1:100) was allowed to generate early glycoforms (**Figure 3A** “Start” panel), the addition of Glc-HMW1ct significantly impacted the resulting product profile (**Figure 3A** “Distraction” panel). Whereas the control reaction quickly proceeded to produce a broad distribution of intermediate glycoforms at low levels (1-Glc to 5-Glc), the distracted reaction revealed the accumulation of 2-Glc as the major product. This change in product profile suggests that Glc-HMW1ct halts the processive phase already at the production of 2-Glc, and enforces the switch to a more distributive mechanism. When the HiNGT-HMW1ct reaction (ratio 1:10) was allowed to form early glycoforms (**Figure 3B**, “Start” panel), the addition of Glc-HMW1ct similarly resulted in the build-up of 2-Glc and 3-Glc as the major products (**Figure 3B**).

**Figure 3.**
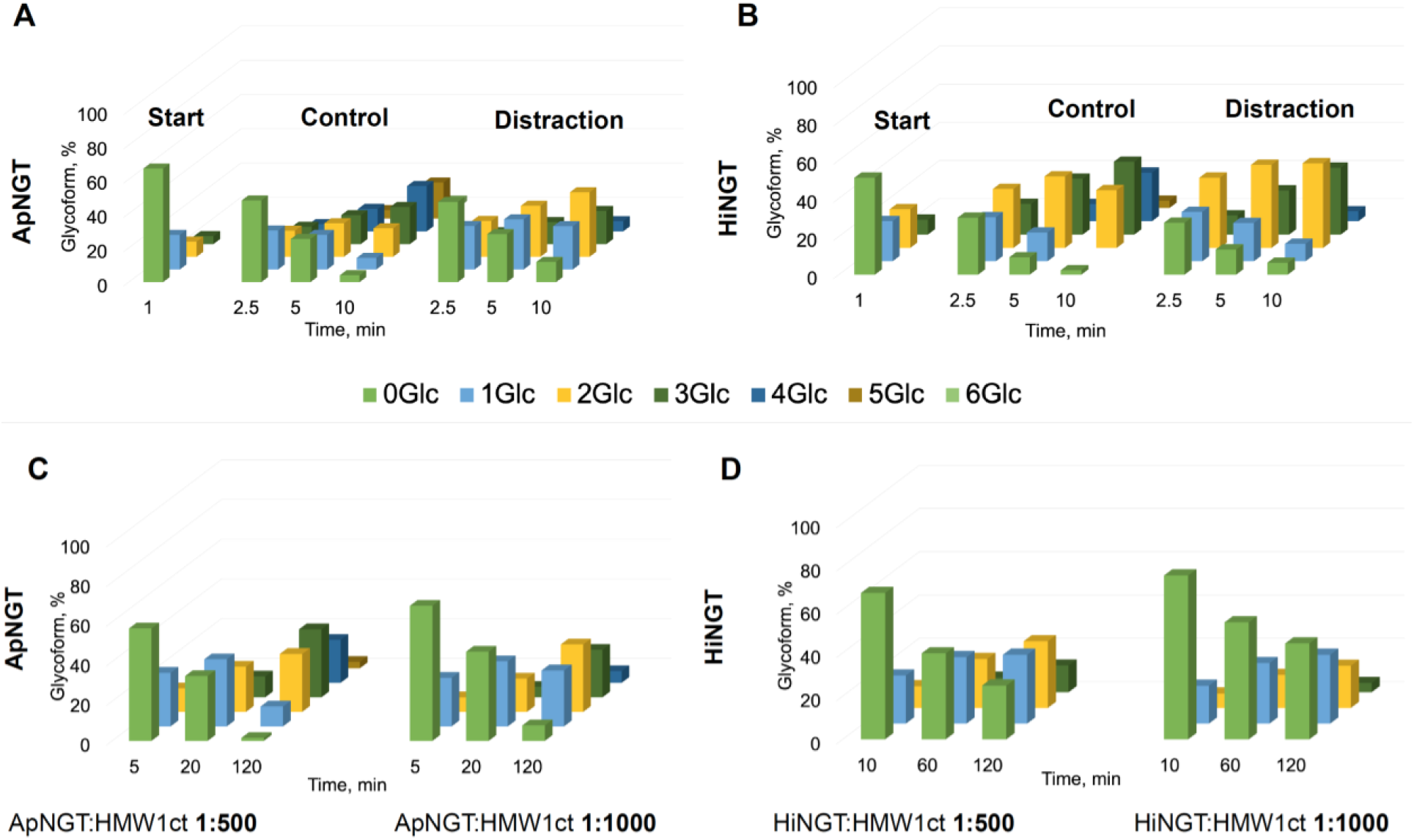
Distraction and single-hit experiments. A) ApNGT:HMW1ct (1:100) was reacted for 1 min, followed by the addition of additional 10 μM Glc-HMW1ct; B) HiNGT:HMW1ct (1:10) was reacted for 1 min, followed by the addition of additional 10 μM Glc-HMW1ct; C) Time-course experiments with ApNGT:HMW1ct at a ratio of 1:500 and 1:1000; D) Time-course experiments with HiNGT:HMW1ct at a ratio of 1:500 and 1:1000; Representative data of two independent experiments are shown. Deconvolved mass spectra for selected time-points are available in Supplementary **Figure S11**.

Although the glycosylated product is able to prematurely halt the processive phase, still a mixture of early glycoforms is persistently produced. This suggests that the early glycoforms (1-Glc to 3-Glc) have an even higher affinity for the NGTs than both non-modified HMW1ct and Glc-HMW1ct. The fast processive phase may be the result of the high affinity for the early glycoforms, which results in a rate enhancement in the early phases of the reaction. To test this hypothesis, an experiment was performed wherein the overnight reaction, containing mostly late glycoforms and showing only very slow glycosylation, was restarted by addition of non-glycosylated substrate (0-Glc) or early glycoforms (0-Glc to 3-Glc).

When the reaction was restarted by the addition of early glycoforms (a mixture of 0,1,2,3,4-times glycosylated HMW1ct, **Figure S12C**) we were intrigued to observe that the reaction proceeded at an increased rate compared to the reaction where non-modified substrate was added (**Figure S12A**), producing late glycoforms in significantly shorter times as compared to the addition of non-modified substrate only. Interestingly, in the case of HiNGT a similar trend was observed (**Figures S12B, S12D**). These results corroborate the findings above that both ApNGT and HiNGT display processive characteristics in the beginning of the reaction.

### Processivity remains under single-hit conditions

As apparent from the initial time-course experiments (**Figure 2**), the observation of processive behaviour seems influenced by the ratio of enzyme to substrate. To understand the impact of the ratio between NGT and HMW1ct, we screened several ratios of both components in a so-called ‘single-hit’ experiment. Characteristic of a single-hit experiment is that the conditions are selected such that multiple binding events are minimized.^26,32^ Generally, this is accomplished with a large substrate-to-enzyme ratio, in which case products bearing multiple modifications can only arise from persistent binding between enzyme and product. In addition, we decided to perform these reactions under dilute conditions, to minimize inhibitory interference by the glycosylated products. **Figure 3C, D** shows the glycoform profiles when HMW1ct was used in large excess to both ApNGT and HiNGT, resulting in enzyme:substrate ratios of 1:500 and 1:1000. Gratifyingly, in all cases the production of early glycoforms (1-Glc to 5-Glc) is apparent, which supports complex formation between NGT and HMW1ct during the first rounds of catalysis. In addition, after overnight incubation the enzymes were inhibited prematurely, generating mixtures of 2-Glc to 5-Glc in the case of ApNGT and 0-Glc to 4-Glc for HiNGT (**Figures S13 and S14**) highlighting the switch from the processive formation of early glycoforms to the subsequent distributive modifications, which are prevented under these single-hit conditions.

### ApNGT and HiNGT prefer glycosylation sites in exposed loops

Having established that ApNGT, and to a lesser extent HiNGT, displays processive characteristics in the initial fast phase, we wondered if NGTs in the fast phase prefer specific sites on HMW1ct. To this end, a site-preference experiment was performed in which the occupancy at all possible sites in HMW1ct was mapped by tryptic digest and LC-MS/MS at early time points. As illustrated in **Figure 4A**, ApNGT preferentially modifies site 9_NAT first (within the first 0.5 minute of the reaction), leading to significant accumulation of the doubly glycosylated peptide (8_NHT+9_NAT), whereas sole modification of site 8_NHT was not observed. This suggests that sites 8 and 9 are modified in a processive manner, without dissociation of the enzyme between the two glycosylation events. Interestingly, also non-sequon site 5’_NAA was modified, which is situated in close proximity to sites 8 and 9, as visualized using a structural model of HMW1ct (**Figure 4C, Figure S1**).^33,34^ After 2.5 minutes, especially di-hexose formation at site 9_NAT appeared (**Figure S15A**). The site preference experiment of HiNGT (at 0.5 minute) reveals a similar preference for site 9_NAT, and this site was also observed with the di-hexose modification (**Figure 4B**). Non-sequon sites 2’_NAG and 9’_NAN were also modified, including with a di-hexose in the latter case. After 20 minutes, modification of sites 5_NVT and 6_NTT appeared, next to di-hexose formation at site 2_NVT and 9_NAT (**Figure S15B**).

**Figure 4.**
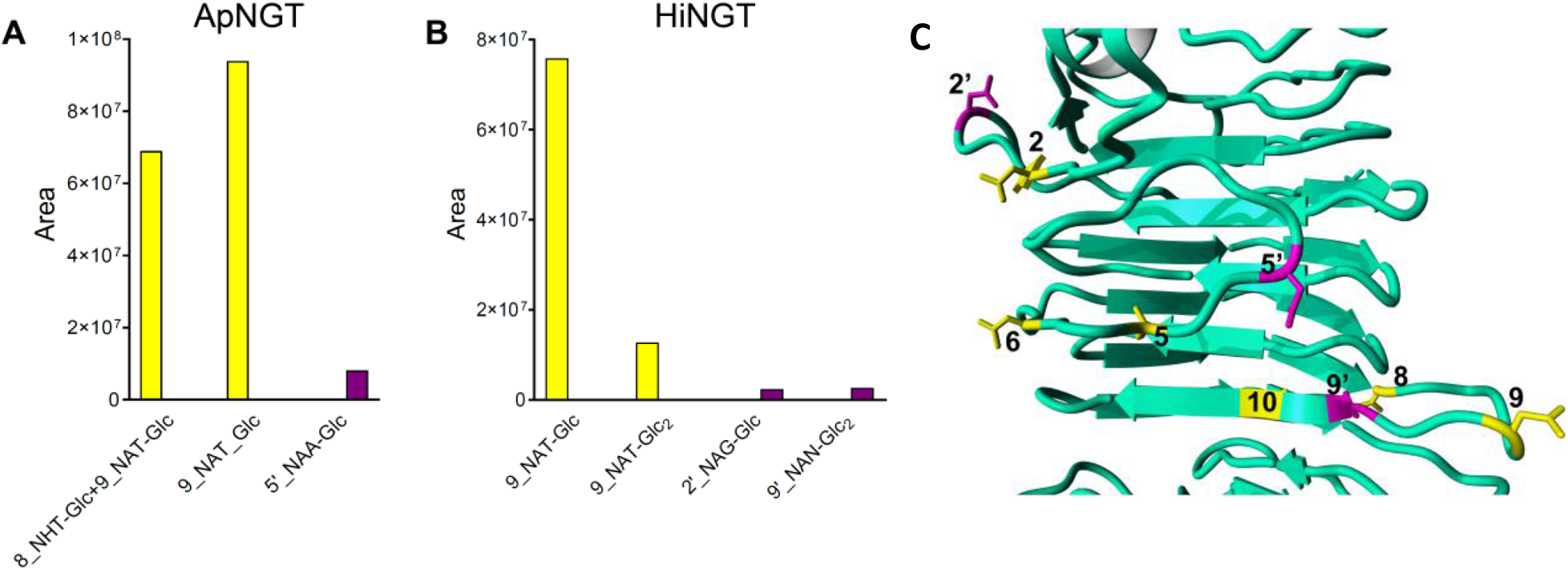
Preference for *N*-glycosylation sites in HMW1ct. A) Site-specific modification for ApNGT after 0.5 min; B) Site-specific modification for HiNGT after 0.5 min; C) I-TASSER model of HMW1ct with sequon sites (yellow) and non-sequon sites (magenta). Representative MS spectra for specific glycosylated peptides are included in **Figures S16-S19**.

The model suggests that HMW1ct adopts an overall β-helix fold, which is a common architecture in bacterial autotransporter passenger domains,^20^ and that all preferred sites are located on exposed loops (**Figure 4C**). Interestingly, although 8_NHT and 9_NAT are located in close proximity, 2_NVT and 5_NVT are situated on the other side of the HMW1ct structure. In addition, both NGTs exhibit some degree of “off-target” glycosylation, in which asparagine residues in non-canonical sequons are modified. Interestingly, these non-sequon sites are predominantly located in the close proximity to the preferred sequon sites (**Figure 4C**) suggesting that when the enzyme is already associated, proximity will drive processive modifications. The di-hexose modification may appear as a result of this proximity-induced binding, however mechanistic insight on the *O*-glycosylation step, as performed by the *N*-glycosyltransferase, is currently lacking.

### ApNGT has a solvent-exposed and relaxed acceptor binding site

Many structural motifs have been associated with processivity, including an extended acceptor binding site, a deep acceptor groove, a closing mechanism with part of the enzyme functioning as a lid, and a ruler helix to control product length.^22,35^ Since there is no precedence for processive character in monomeric protein glycosyltransferases, we set out to identify the possible structural elements that are responsible for processivity using docking and molecular dynamics (MD) simulations. We selected ApNGT because there is one report of a crystal structure with UDP bound (PDB: 3Q3H).^36^ First the glucose was added to generate a docked structure of ApNGT::UDP-Glc, which was used as a scaffold for peptide docking. The similarities between hOGT and ApNGT are evident when comparing UDP-GlcNAc and UDP-Glc, respectively (**Figure 5A**), to nucleotide-sugar conformations from several other complexes within the GT-B enzyme family (*i*.*e*. inverting enzymes MurG, UGT71G1, UGT72B1, VvGT1, and retaining enzymes AGT, OtsA, WaaG).^37^ The unusual UDP-sugar pyrophosphate conformation positions the α-phosphate to act as the proton acceptor in the hOGT-catalyzed glycosylation reaction.^37^ In this regard, the pyrophosphate torsion angles of UDP-Glc are more similar to the angles of UDP-GlcNAc in hOGT than to the angles of all the other nucleotide-sugar structures. Protein-ligand interactions in the UDP-sugar binding site resemble those observed in hOGT (**Figure S20**).

**Figure 5.**
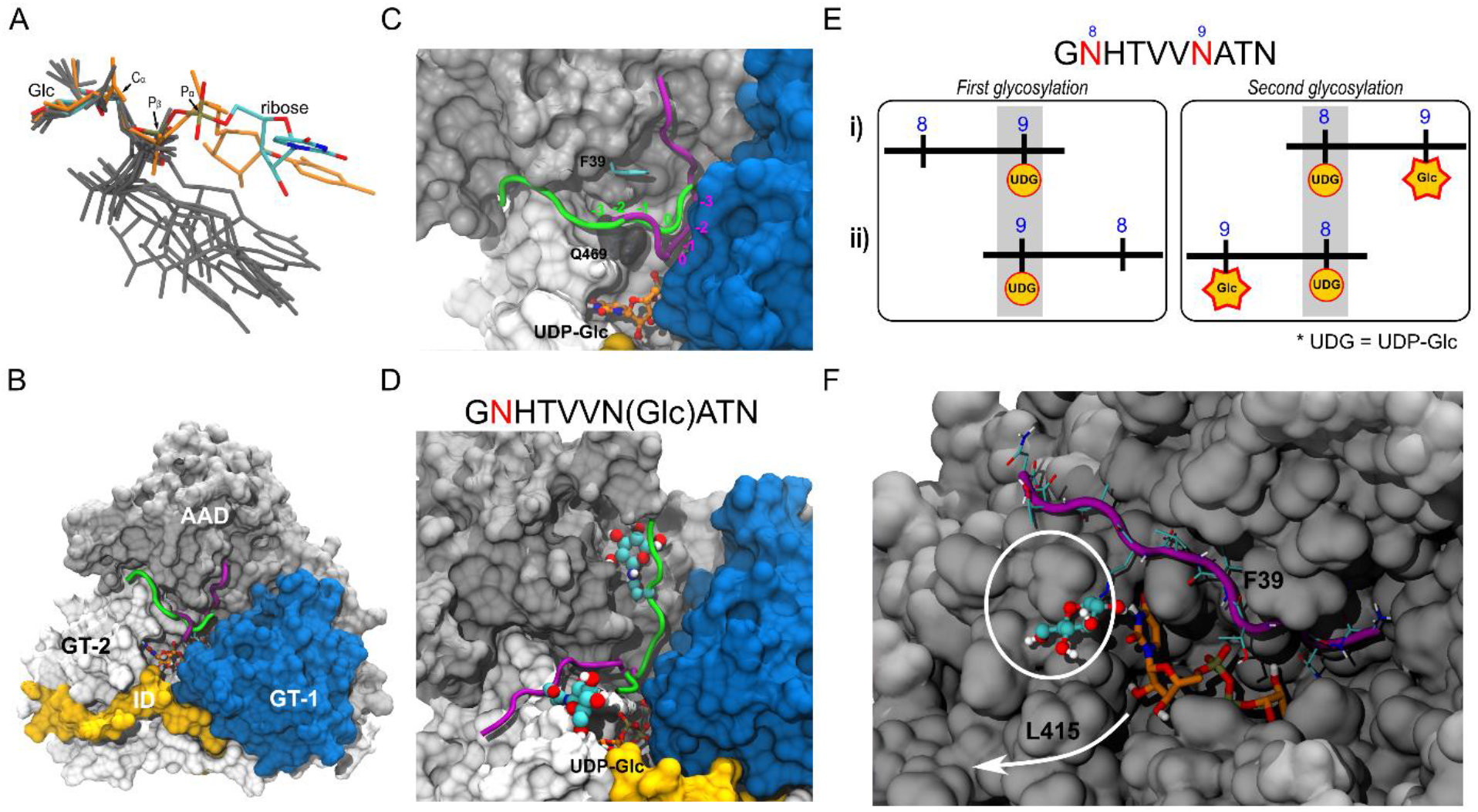
Docking and MD simulation of the ApNGT::peptide::UDP-Glc complex reveals relaxed acceptor binding. A) The pyrophosphate torsion angles UDP-Glc in ApNGT (colored sticks) are more similar to the pyrophosphate angles of UDP-GlcNAc in hOGT (orange sticks), than to other glycosyltransferases in the GT-B family (grey sticks). B) Two binding modes of peptide G**<u>N</u>**(8)HTVV**<u>N</u>**(9)ATN in ApNGT found by computational modeling presented in the purple and green cartoons (opposite N→C directions). Both peptides are bound to UDP-Glc by Asn(9) (shown in stick); C) Close-up structure of the binding modes for the peptide G**<u>N</u>**(8)HTVV**<u>N</u>**(9)ATN, docked to UDP-Glc via Asn(9); D) Close-up structure of the binding modes for the peptide G**<u>N</u>**(8)HTVV**<u>N</u>**(Glc)ATN, docked to UDP-Glc via Asn(8); E) Schematic representation of the possible mechanisms in which the peptide is docked at site Asn(9), is glycosylated, and then slides in the forward direction (i) to allow glycosylation at site Asn(8), or in the reversed direction (ii); F) Space-filling model of the ApNGT::Glc-peptide::UDP-Glc complex that suggests there is enough space for UDP to dissociate and UDP-Glc to associate in between glycosylation events.

Next, the complex of ApNGT::UDP-Glc with the peptide GN(8)HTVVN(9)ATN (corresponding to HMW1ct sequons 8 and 9) was created to assess possible binding poses of the preferred adhesin fragment (**Figure 5B**). The nucleophilic N from Asn(9) was constrained to be in close proximity to the anomeric C_α_ carbon, and peptide binding modes were generated. The binding site of ApNGT was found to be flexible enough to allow several peptide binding modes (**Figure 5C**, main binding modes in green and purple) near the postulated acceptor binding groove and making contacts with the proposed acceptor binding residues Phe39, His272, His277, and Gln469.^36,38^ Our results suggest that the peptide-binding region in ApNGT is located on the solvent-exposed enzyme surface. In contrast, in hOGT the unfolded peptide binds in a groove that is located inside a superspiral formed by repeated TPR regions.^39^ The known crystal structures of hOGT show two binding modes either with a shallow pose (**Figure S21**, purple cartoon) or more embedded pose in the TPR domain (**Figure S21**, green cartoon), where the former recognizes semi-folded peptide regions, and the latter is for extended peptides.^39^ Interestingly, ApNGT revealed unexpected flexibility in peptide binding, and opposing orientations with respect to the *N*- and *C*-termini appeared to bind stably (**Figure 5C**). In contrast, the crystal structures of hOGT show the peptides in only one orientation (**Figure S21**).

As the experimental data suggests that one Glc modification promotes a second Glc-transfer, we generated peptide-enzyme complexes with the glycosylated peptide G**<u>N</u>**(8)HTVV**<u>N</u>**(Glc)ATN, with preferred site 9_NAT glycosylated (*vide supra*). Two regions for the binding of the Glc moiety were found (**Figure 5D**, space-filling models), but none of these displayed increased affinities. Interestingly, the Interface Score of the peptide-protein complex, with and without glycosylation, was around −35 kcal/mol, suggesting similar binding energies for both peptide and Glc-peptide. MD simulations of the Glc-peptide complex did not show Glc-focused interactions with ApNGT. Based on the computational modeling, we hypothesize that after glycosylation of the first site (N(9)AT), the peptide slides along the enzyme to achieve a second glycosylation at N(8)HT, while anchoring to the enzyme with its N(Glc)AT site (**Figure 5E**). Because ApNGT is flexible in the N-to C-terminus direction the peptide binds, this process could potentially happen in the opposite direction. In addition, the model suggests enough space for UDP to dissociate and be substituted for a new UDP-Glc, to continue catalysis (**Figure 5F**).

## Discussion

Protein glycosyltransferases are abundantly present in all domains of life, and are found to catalyse a wide range of protein modifications, with new examples emerging at a steady pace.^40^ They show an intriguing level of diversity in specificity for both sugar donors and protein substrates, but also recognition elements (amino acid residues, structural folds) and timing of modification (co- or post-translational). As protein glycosylation is not genetically encoded, the spatiotemporal drivers and effects of protein glycosylation are at the same time exciting and challenging to study.

Our results reveal how ApNGT, and to a lesser extent HiNGT, perform hyperglycosylation of HMW1ct adhesin in a two-phase mechanism (**Figure 6**). In the beginning of the reaction, ApNGT glycosylates HMW1ct using a processive mechanism that yields a broad distribution of intermediate glycoforms. Compared to the starting substrate HMW1ct, especially the early glycoforms seem to be suitable substrates for processive modification, which is a characteristic of processive enzymes. However, the enzyme-substrate complex is receptive to the presence of the fully modified Glc-HMW1ct product that successfully competes with binding to the enzyme, resulting in a shortening of the processive phase. After this fast processive phase, both ApNGT and HiNGT are increasingly inhibited by the high affinity for the glycosylated product Glc-HMW1ct, and only incrementally add glucose residues to remaining sites. The fact that di-hexose formation and modification of non-sequon sites generally happens on and in close proximity to defined sequons further strengthens the hypothesis that NGTs employ proximity-induced processive glycosylation. However, whether NGTs stay fully associated to ensure processivity, or that they engage in ‘hopping’ (*i*.*e*. microscopic dissociation followed by quick reassociation), in analogy to processivity in DNA-binding proteins, is currently impossible to determine.^41,42^ A hallmark of processivity is the high affinity of the enzyme for its product. Therefore, processive enzymes may be more sensitive to product inhibition than enzymes that employ a distributive mechanism.^43^ Conversely, because distributive enzymes dissociate after catalysis, they may also be susceptible to competitor binding. For distributive protein kinases, an increase in substrate concentration results in accumulation of partially phosphorylated species, that serve as competitive kinase inhibitors.^30^ As the NGTs studied here display characteristics of both processes, we suggest to denote the mechanism of these NGTs as semi-processive. We propose a mechanistic model that starts with NGT binding to HMW1ct, followed by fast and processive glycosylation of adjacent sites facilitated by sliding over the NGT surface (**Figure 6A**), or di-hexose formation (**Figure 6B**). We expect that this promiscuous surface-binding is a structural basis for processivity, as the lack thereof may be at the basis of the distributive character observed in hOGT.^23,36^After a few additional modifications, NGT enters a slower distributive phase, in which it may randomly bind to both sequon and non-sequon sites on the surface of HMW1ct. The resulting products have high affinity for the NGTs, resulting in retardation of glycosylation by product inhibition. Together, this leads us to propose a semi-processive mechanism for NGTs.

**Figure 6.**
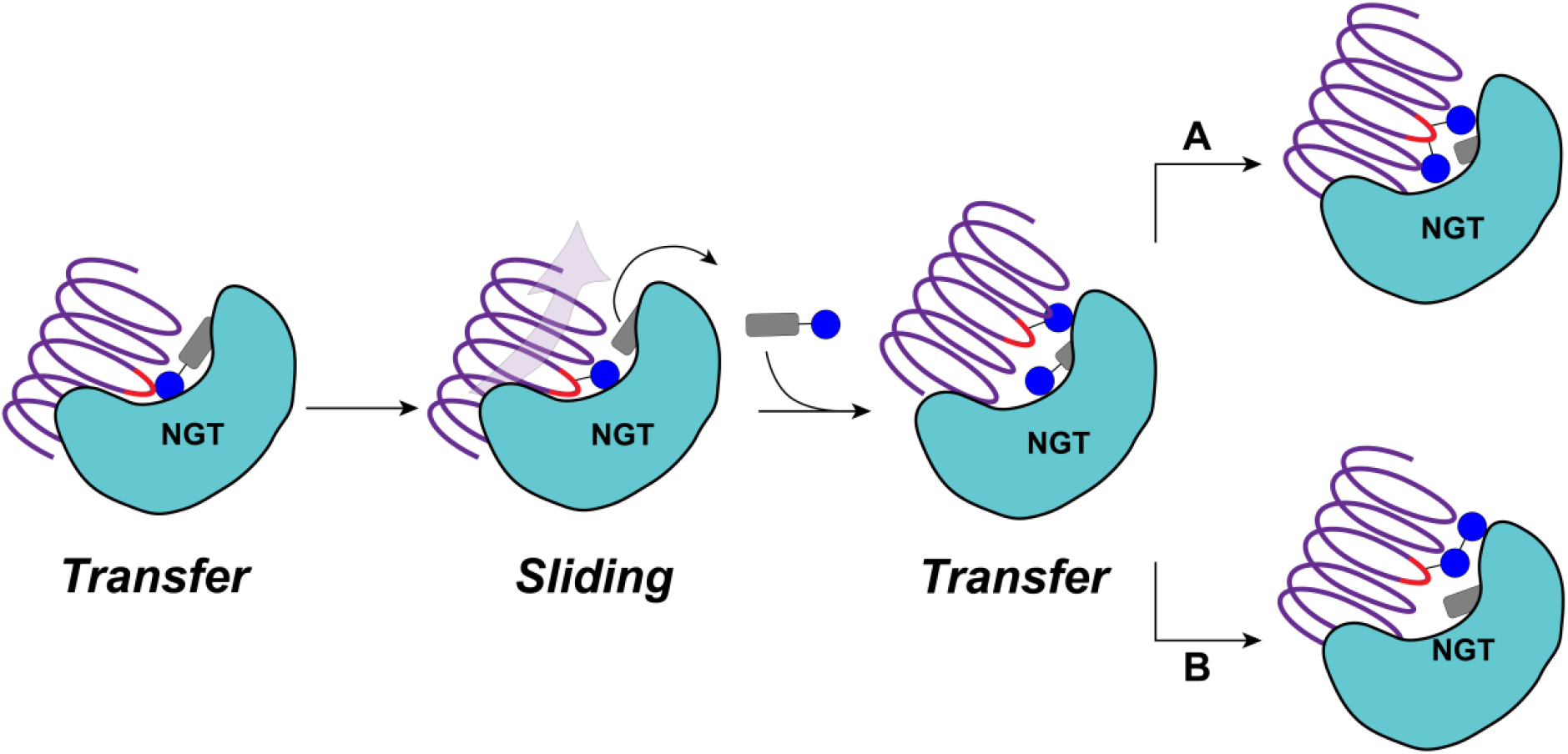
Model for the sliding mechanism in the fast phase in the semi-processive glycosylation of HMW1ct by NGTs, that results in processive glycosylation of adjacent sites (A), and di-hexose formation (B).

NGTs have a high preference for sequons that are exposed on the surface of the acceptor protein, which is consistent with the post-translational timing of the modification.

Moreover, especially the bacterial adhesins and autotransporters share a general β-helical fold,^44,45^ which is also highly associated with two-partner secretion proteins in different species.^46,47^ It will be highly revealing to investigate other known and predicted NGTs for processive characteristics,^48^ and revisit currently known β-helical adhesins to find an associated NGT.

The clear processive features in the NGTs under study here raises the question of the functional relevance. Processivity is well-established in template-driven production of oligonucleotides. For post-translational modifications, such as phosphorylation and glycosylation, there is little knowledge on the importance of multisite modifications, but the sheer number of modifications may seem more important than the specific locations. The high association rate of the substrate and processivity of early glycoforms may ensure a high level of Glc-modifications on the HMW adhesins before export by the HMW1B translocator. In general, the density of epitopes is directly linked to the efficiency of natural multivalent interactions, and is proposed to serve as a mechanism to regulate the biological interaction.^49^ Multisite glycosylation may be an elegant solution to ensure efficient bacterial attachment to receptors through multivalency,^50-52^ to overcome the generally poor (mM range) affinity of proteins for carbohydrate ligands.

The knowledge that NGTs can support processive characteristics is important in the biotechnological use of such enzymes to create well-defined glycoproteins. Several studies have focused on employing NGTs (and their engineered variants) in the biosynthesis of defined glycoproteins for biotechnological applications and vaccine development.^38,48,53-55^ The ApNGT mutant Q469A showed reduced product inhibition, and produced a more homogenously glycosylated HMW1ct, with up to 10 residues. Based on the central position of Q469 in both UDP-Glc and peptide binding as revealed by molecular modeling, we propose that Q469 may function as a ‘processive switch’, preventing the glycosylated product from leaving the binding site, and thereby increasing the association required for an additional round of catalysis.^38^ Sequence alignment indicates a corresponding Gln residue in a conserved region in HiNGT (Gln495, **Figure S22**), but without more structural information, it is difficult to assess its involvement in the mechanism.

Our results suggest that glycoprotein production systems based on NGT expression in *E. coli* may suffer from the low UDP-Glc levels (typically, 1-2 mM),^56^ as that may lead to premature product inhibition. In agreement with other reports,^31^ we found that the glycosylation of HMW1ct is highly dependent on the levels of NGTs. This may limit the usefulness of future vaccines against glycosylated HMW adhesins, as bacteria may express different NGT levels under changing conditions, resulting in differently glycosylated adhesins. Moreover, glycosylated HMW1ct has been linked to the production of pathogenic auto-antibodies in multiple sclerosis, raising the question whether glycosylated HMW1 adhesin are in general appropriate vaccine targets.^21^ We show here that processivity in NGTs arises from high affinity for the intermediate products, and this may inspire a class of inhibitors that capitalize on product binding, for instance by generating glycosylated β-helical peptide scaffolds.

In summary, we have provided evidence that both ApNGT and HiNGT display processive characteristics in the first fast phase of HMW1ct glycosylation, followed by a phase with distributive features, together resulting in a semi-processive mechanism. Molecular modeling reveals that ApNGT has promiscuous substrate binding preference, which allows for sliding of the enzyme along the adhesin surface. Further investigations into the mechanisms of other bacterial NGTs will reveal whether processivity is a general mechanism that bacteria use to achieve hyperglycosylation of extracellular proteins involved in virulence.

## Methods

### Protein expression and purification

HiNGT-His_6_ was generated from the HMW1C gene extracted from the genomic plasmid isolated from *H. influenzae* R2846 (provided by Prof. Arnold Smith) and incorporated in a pET24 plasmid. The genes encoding His_6_-HMW1ct (pET45) and ApNGT-His_6_ (pET24) were overexpressed and proteins were purified as described.^21^ In short, pET24 plasmid encoding ApNGT or HiNGT (or pET45b plasmid encoding HMW1ct) was transformed into BL21 DE3 *E. coli* cells by heat shock and plated onto Luria-Bertani agar plates with 100 µg/mL kanamycin (or ampicillin for HMW1ct). A 10 mL starter culture (with 100 µg/mL of appropriate antibiotic) was prepared and grown overnight at 37 °C with shaking. A portion of the starter culture was used to inoculate a larger volume of TB (Terrific Broth) (0.5 L with 100 µg/mL of appropriate antibiotic), which was incubated at 37 °C with shaking until optical density readings reached 0.6-0.8. At this point protein overexpression was induced by addition of 1 mM final concentration isopropyl β-thiogalactopyranoside (IPTG) and incubated at 16°C with shaking overnight. Cells were harvested by centrifugation (5000 rpm), resuspended in ice-cold lysis buffer (50 mM HEPES, 100 mM NaCl, 10% glycerol, pH 7.5) and lysed by sonication (Branson Sonifier 450: 30% duty cycle, 2.5min) in the presence of the protease inhibitors cocktail (Roche). Cell debris was removed by centrifugation and supernatant was used for Ni-affinity chromatography purification. Briefly, 3-4 mL of Ni-NTA resins (Qiagen) were applied on the gravity column, washed with water and equilibrated with the lysis buffer. Cell-free extract was then mixed with the resins for 1.5 h at 4°C with gentle shaking. Afterwards, cell-free extract was allowed to flow through and resin-bound proteins were washed twice with washing buffer (50 mM HEPES, 300 mM NaCl, 5% glycerol, 15 mM imidazole, pH 7.5) and then eluted in three steps with elution buffer (50 mM HEPES, 300 mM NaCl, 5% glycerol, 400 mM imidazole, pH 7.5). Fractions containing protein of interest were collected and desalted using PD-10 midi desalting columns (GE Healthcare). Protein of interest was eluted with the storage buffer (50 mM HEPES, 100 mM NaCl, 10% glycerol, pH 7.5), aliquoted and stored at −80 until further use.

### Glc-HMW1ct production by protein co-expression, purification and AE separation

A pET24 plasmid encoding ApNGT-His_6_ and pET45 plasmid encoding His_6_-HMW1ct were simultaneously transformed into *E. coli* BL21 DE3 by heat shock and plated onto Luria-Bertani agar plates with 100 µg/mL of both kanamycin and ampicillin. Culture growth, protein co-overexpression and purification were carried out in the same way as described in the previous section. The separation of ApNGT and glycosylated HMW1ct was performed via anion exchange on FPLC ÄKTA Pure system. The protein mixture was first desalted and eluted with FPLC buffer A (20 mM Tris, 20 mM NaCl, pH 8) and then in 2 mL injections applied to an anion exchange column (HiTrap, Q FF, 5 mL). A gradient of 0 to 50% of buffer B (20 mM Tris, 1 M NaCl, pH 8) was applied with the constant system flow of 5mL/min which resulted in a baseline separation of glycosylated HMW1ct and ApNGT. Fractions containing Glc-HMW1ct were collected and concentrated with Amicon spin filter columns and high-salt content of the elution buffer was removed via diafiltration with the storage buffer (50 mM HEPES, 100 mM NaCl, 10% glycerol, pH 7.5).

### *In vitro* glycosylation of HMW1ct: time course of glycosylation, various ratios

To monitor the time-course of HMW1ct glycosylation by ApNGT, reaction mixtures contained final concentrations of 10 µM HMW1ct, 1mM UDP-Glc and 0.1 µM ApNGT (for 1:100 ratio) or 1 µM ApNGT (for 1:10 ratio) in buffer (50 mM HEPES, 100 mM NaCl, 10% glycerol, pH 7.5). To monitor the time-course of HMW1ct glycosylation by HiNGT, reaction mixtures contained final concentrations of 10 µM HMW1ct, 1 mM UDP-Glc and 0.1 µM HiNGT (for 1:100 ratio) or 1 µM HiNGT (for 1:10 ratio) in buffer (50 mM HEPES, 100 mM NaCl, 10% glycerol, pH 7.5). Aliquots of 50 µL were taken after 1min, 5min, 10min, 30min, 60min, 120min, 300min, 960min, quenched with 50 µL boiling water and incubated at 100 °C for 10min.

### Intact protein MS data deconvolution and quantification

Samples of quenched reaction aliquots were further diluted two-fold with ultrapure water to reduce the viscosity and subjected to intact protein LC-MS analysis. The runs were performed on a Thermo Ultimate 3000 QExactive Orbitrap instrument (Thermo Scientific) or Orbitrap Velos Pro instrument (Thermo Scientific) equipped with a C8 column (Aeres C8, 150×2.1 mm, 3.6 µm). Injection volume was 1 µL. The gradient started at 25% B (ACN, 0.1% FA) and was increased to 90% in 10 min, where it stayed for 2 min with subsequent decrease back to 25% in 3min. The flow was 0.35 mL/min and the column temperature were kept at 60 °C. The deconvolution was performed using open access UniDec software, where the raw spectrum of charged states was exported and settings outlined in **Table S3** were applied. Next, protein glycoforms (**Table S4**) and their intensities were exported into Excel for analysis and construction of graphs. Intensities of all glycoforms at a certain time point were summed to obtain total protein count at that point. The intensities were then corrected for the largest amount of total protein and converted into percentage value. These values were then plotted in 2D graphs. As an additional control of protein ionization differences, the intensities of 10 µM of non-glycosylated HMW1ct and fully glycosylated Glc-HMW1ct were compared, as was a mixture of both components at 10 µM (**Figure S23**). Whereas the separate components showed similar ionization intensities, the mixed sample revealed reduced intensity for Glc-HMW1ct. Since we are interpreting the results based on product profiles instead of absolute intensities, we decided to not introduce a correction factor for ionization intensity.

### Continuous coupled assay

Spectrophotometric assays were performed in 96-well plates (total volume per well 140µL) containing 50 mM HEPES, 100 mM NaCl, 25 mM MgCl_2_, 300 units pyruvate kinase, 20 units lactate dehydrogenase, 250 µM NADH, 500 µM phosphoenolpyruvate and 0.1 µM of NGT (for 1:100 ratio) or 1 µM NGT (for 1:10 ratio). Absorbance at 340 nm was monitored over time in the BioTek Synergy H1 plate reader until a steady baseline was reached (around 3 min). Subsequently, UDP-Glc was added to a final concentration of 1 mM and the baseline was monitored for another 3 min. Reactions were initiated by addition of HMW1ct to a final concentration of 10 µM and the plate was incubated in the plate reader at 25 °C. All experiments were performed in triplicate (time-course experiments) or duplicate (kinetic parameter determination). For the determination of kinetic parameters, the NGT concentration was kept at 0.1 µM and a range of HMW1ct concentrations was used: 1 µM, 2.5 µM, 5 µM, 10 µM, 25 µM, 50 µM, 75 µM. UDP-Glc concentration was kept at 5 mM. Steady-state rates (*V*_0_) were calculated from the slope of the linear portion of the decline in absorbance over time calculated with ε = 6,300 M^-1^cm^-1^. Non-linear regression and preparation of graphs was performed with GraphPad.

### Restarting overnight reaction: substrate and early glycoforms

For this experiment, an overnight reaction with 1:50 ratio (ApNGT-HMW1ct) or 1:5 ratio (HiNGT-HMW1ct) was prepared in buffer (50 mM HEPES, 100 mM NaCl, 10% glycerol, pH 7.5). Briefly, 200 µL reaction mixtures contained 0.2 µM ApNGT (or 2 µM HiNGT), 10 µM HMW1ct and 1 mM UDP-Glc. Overnight reaction was split into three (45 µL) and equal volume of a) buffer; or b) 20 µM HMW1ct (final concentration 10 µM) or c) 20 µM early glycoforms (final concentration 10 µM) were added. As a result, a typical 1:100 ratio (ApNGT-HMW1ct) or 1:10 ratio (HiNGT-HMW1ct) was reached. Early glycoforms were generated by incubating 20 µM HMW1ct with 0.2 µM ApNGT and 1mM UDP-Glc for 1.5 min in buffer (50 mM HEPES, 100 mM NaCl, 10% glycerol, pH 7.5), and then quenching the reaction by incubation at 100°C for 10min. For quenching, 50 µL aliquots were taken after 1, 5, 10 and 30min, mixed with equal volume boiling water and incubated at 100°C for another 10 min.

### Distraction assay with (Glc)HMW1ct

The typical glycosylation reactions were prepared with 1:100 ratio (ApNGT-HMW1ct) or 1:10 ratio (HiNGT-HMW1ct). Generally, 500 µL total volume contained final concentration of 0.1 µM of ApNGT or 1 µM of HiNGT, 10 µM of HMW1ct and 1 mM of UDP-Glc in buffer (50 mM HEPES, 100 mM NaCl, 10% glycerol, pH 7.5). Reactions were allowed to proceed for 1 min, at which point an aliquot was taken and quenched as described above. The rest of the reaction mixtures was split into two (200 µL each) and either 28.5 µL of buffer (control) or 28.5 µL of 80 µM Glc-HMW1ct (distraction, final concentration 10 µM) were added to the 200 µL of the ongoing glycosylation reaction. Aliquots of 50 µL were taken after 1.5 min, 4 min and 9 min and quenched as described above.

### Single hit conditions

To prepare a 1:500 enzyme-to-substrate ratio reaction, a total volume of 200 µL (or 600 µL for a longer time-course experiment) contained 0.1 µM of NGT enzyme, 50 µM of HMW1ct and 1mM UDP-Glc in buffer (50 mM HEPES, 100 mM NaCl, 10% glycerol, pH 7.5). For 1:1000 ratio, 300 µL reaction mixtures were prepared with 0.065 µM enzyme, 65 µM HMW1ct and 1mM UDP-Glc in buffer (50 mM HEPES, 100 mM NaCl, 10% glycerol, pH 7.5). Aliquots of 50 µL were taken at 5 min, 20 min, 120 min (for ApNGT-HMW1ct), and 10 min, 60 min, 120 min (for HiNGT-HMW1ct) and quenched as described above. For the longer time-course experiments, aliquots were takes after 1 min, 5 min, 10 min, 30 min, 60 min, 120 min, 180 min and 960 min.

### Same ratio, different concentrations

For both ApNGT and HiNGT, four reaction mixtures were prepared in buffer (50 mM HEPES, 100 mM NaCl, 10% glycerol, pH 7.5), wherein enzyme and substrate concentrations were varied to achieve 1:100 ratio. Briefly, for ApNGT-HMW1ct reaction mixtures of 50-100 µL contained final concentrations of a) 0.05 µM ApNGT and 5 µM HMW1ct; b) 0.1 µM ApNGT and 10 µM HMW1ct; c) 0.5 µM ApNGT and 50 µM HMW1ct; d) 1 µM ApNGT and 100 µM HMW1ct. The UDP-Glc concentration was either kept at 1mM, or adjusted for each reaction and kept at 100-fold excess over HMW1ct (1 mM for a and b, 5 mM for c and 10 mM for d). HiNGT-HMW1ct reaction mixtures of 50-100 µL contained final concentrations of a) 0.5 µM HiNGT and 5 µM HMW1ct; b) 1 µM HiNGT and 10 µM HMW1ct; c) 5 µM HiNGT and 50 µM HMW1ct; d) 10 µM HiNGT and 100 µM HMW1ct. All reactions were incubated overnight at room temperature and 50 µL aliquots were taken the next day and quenched as described above.

### Site preference investigation

The site preference of NGTs was determined via proteomics analysis with subsequent data quantification. For the glycosylation site preference two experiments were ran in parallel: a glycosylation reaction (1:100 ratio for ApNGT and 1:10 for HiNGT) and a blank reaction (all components except UDP-Glc) to achieve comparable protein levels for later quantification purposes. Briefly, for ApNGT-HMW1ct total volume of 350 µL contained 0.1 µM ApNGT, 10 µM of HMW1ct and 1 mM of UDP-Glc (or none for the blank) in buffer (50 mM HEPES, 100 mM NaCl, 10% glycerol, pH 7.5). For HiNGT-HMW1ct, the total volume of 100 µL contained 1 µM HiNGT, 10 µM HMW1ct and 1mM UDP-Glc (or none for the blank) in buffer. The aliquots were drawn at early time points for both enzymes (ApNGT: 0.5 min and 2.5 min, HiNGT: 0.5 min, and 20 min) when presumably only preferred glycosylation sites are being modified. The reaction aliquots were quenched as described above and then subjected to trypsin digestion and proteomics analysis, as described below.

### Proteomics data quantification and analysis

From the purified protein sample, 100 µg was taken and 100 mM ammonium bicarbonate (ABC) was added to reach 32 µl. 8 µL 8 M urea was added to obtain a concentration of 1.6 M urea. Furthermore, 1 µl of 0.2 M TECEP was added. The sample wad mixed and incubated at 37 °C for 1 h. After the incubation, the sample was cooled to room temperature. Alkylation of cysteines was performed by adding 1 µL of freshly prepared 0.4 M iodacetamide and incubated at 25°C for 30 min in the dark. The pH was checked to be around 8-9 and if required adjusted using 1 M ABC. Trypsin (Promega, V5113) was added at a ratio of 1:50 w/w trypsin/protein and incubated overnight at 37°C. Sample clean-up by solid phase extraction was performed with Pierce® C18 tips (Thermo, 87784) according to the supplier’s manual. The eluate fraction was dried under vacuum and reconstituted with 20 µL 2% acetonitrile, 0.1% formic acid. Peptide separation was performed with 2 µL peptide sample using a nano-flow chromatography system (EASY nLC II; Thermo) equipped with a reversed phase HPLC column (75 µm, 15 cm) packed in-house with C18 resin (ReproSil-Pur C18–AQ, 3 µm resin; Dr. Maisch) using a linear gradient from 95% solvent A (0.1% FA, 2% acetonitrile) and 5% solvent B (99.9% acetonitrile, 0.1% FA) to 28% solvent B over 45% at a flow rate of 200 nl/min. The peptide and peptide fragment masses were determined by an electrospray ionization mass spectrometer (LTQ-Orbitrap XL; Thermo).

### Data processing

hermo raw files were imported into the Peaks Studio software (Bioinformatics Solutions) analyzed against forward and reverse peptide sequences of the expression host *E. coli K12* and the over-expressed construct HMW1ct. The search criteria were set as follows: specific tryptic specificity was required (cleavage after lysine or arginine residues but not when followed by a proline); three missed cleavages were allowed; carbamidomethylation (C) was set as fixed modification; oxidation (M) and deamination (NQ) as variable modification. Variable glycosylation modification was set to 1, 2 or 3 hexose(s) (N). The mass tolerance was set to 15 ppm for precursor ions and 0.5 Da for the fragment ions.

### Data analysis

Raw data files were processed with PEAKS X Plus software and a search for the glucosylation modification (+162.05 Da; +324.1 Da or 485.15 Da) on asparagines was applied. To obtain the intensities of various (glyco)peptides and perform quantitative analysis, the peptide lists were exported from PEAKS software and merged together to obtain intensities of the peptides of both blank samples and reaction samples at various time points. To analyze which sites were glycosylated first, glycopeptides were sorted by glycosylation sites (1 to 12, **Table S5**) and intensities of respective glycopeptides that contain the sites of interest were summed. These values were plotted against time to show the rise in intensity for the peptides bearing certain glycosylation sites over others (preferred sites). Subsequently, glycosylated peptides were manually inspected to confirm the presence of the signature ions. Spectra that featured insufficient fragmentation patters around the site of glycosylation to conclude site-specific glycosylation were discarded.

### Processivity parameters calculations

Calculations were performed as described before.^25,26^ The percentage of active NGT was calculated using the following formula:

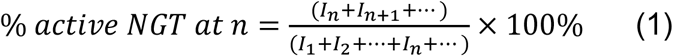

*I*_1_ – intensity of the product modified with 1-Glc; *I*_*n*_– intensity of the product modified by n Glc. Processivity factor was calculated using the formula (2). Formula (3) represents the processivity factor expressed in intensity values.

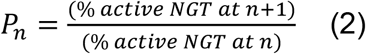

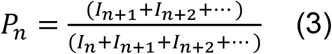

### Affinity studies by surface plasmon resonance

Surface plasmon resonance experiments were performed on a Biacore 3000 instrument (GE Healthcare). Non-glycosylated adhesin HMW1ct (substrate) and glycosylated adhesin Glc-HMW1ct (product) were immobilized to different flow cells on CM5 sensorchip via standard primary amine coupling in 10 mM sodium acetate, pH 5, to densities of c. 5000 response units. To determine the binding affinity for its protein substrate/product in buffer HBS-EP (0.01 M HEPES pH 7.4, 0.15 M NaCl, 3 mM EDTA, 0.005% v/v Surfactant P20) at 25°C, series of increasing concentration of ApNGT (0.125 µM, 0,25 µM, 0.5 µM, 1 µM, 2 µM, 4 µM, 8 µM, 16 µM) were flushed at 50 µL/min for 60 s over the chip with immobilized adhesin proteins after which the dissociation was followed for 4000 s. The SPR signal was corrected for the signal of an empty flow cell, and these data were analysed by BIAevaluation v. 4.1.1 software (GE Healthcare) using a global fitting analysis. Fitting according to the model “heterogeneous ligand” gave an improved fit compared to the 1:1 Langmuir model, indicating that chemical immobilization of the adhesins led to a small fraction of the proteins having a different interaction with the analyte. Kinetic values are averaged over 5 independent binding experiments, using either the full concentration range or a subset of concentrations.

### Molecular modeling

The crystal structure of ApNGT (PDB code: 3Q3H) contained coordinates for the UDP molecule but not for the Glc moiety in UDP-Glc. Thus, we first produced a ApNGT::UDP-Glc docking complex to serve as scaffold for peptide docking. The peptide GNHTVVNATN, corresponding to residue numbers 193 – 201 of the adhesin fragment, were docked into ApNGT::UDP-Glc.

### Docking of UDP-Glucose

The initial geometries for the UDP-Glc ligand were obtained by collecting 100 crystal structures of non-covalently bound UDP-Glc from the PDB databank, belonging to a wide range of enzyme families. The UDP-Glc conformer library was used for docking with the Rosetta (v2020.11) Enzyme Design application. The best structures were selected based on the Rosetta Interface Energy, and a representative ApNGT::UDP-Glc structure served as scaffold for all peptide docking calculations.

### Peptide docking

Peptide docking consisted of three subsequent stages: rigid-body docking, moderate movement refinement, and minimal movement refinement. The first stage was intended to produce a coarse model of the protein-peptide complex. The peptide was treated as a rigid molecule by directly providing a conformer library consisting of 1,000 peptide conformers generated using MODPEP.^57^ A distance constraint of 3.0 ± 0.5 Å (penalty 50.0) between Asn-ND2 and the Cα of UDP-Glc was used to restrict the peptide conformations to poses relevant for catalysis. A total of 10,000 structures from 100 individual seeds were generated, and ranked. Ranking was done based on two criteria: 1) distance between Asn-ND2 and the Cα of UDP-Glc, and 2) the Interface Score of the enzyme-peptide complex. The top 100 ranking structures were selected for refinement. The difference between the moderate movement and the minimal movement refinement protocols is that the former performs centroid-based movements with larger perturbations before the all-atom refinement stage. Both protocols employed the FlexPepDock^58,59^ Rosetta application. In each subsequent refinement stage, a total of 10,000 enzyme-peptide structures were generated, and the selection of the top 100 ranked models consisted of three criteria: 1) the distance between Asn-ND2 and the Cα of UDP-Glc, 2) the Interface Score (*I_sc*) of the enzyme-peptide complex, and 3) the total Rosetta Score (*total_score*). Interestingly, the Interface Score of the peptide-protein complex, with and without glycosylation, was around −35 kcal/mol (*I_sc*), suggesting similar binding energies for both peptide and Glc-peptide. While the glycosylation in GNHTVVNATN did not improve the score, there are two factors to consider when comparing the Interface Score of the glycosylated and non-glycosylated peptide. The Rosetta Energy Function,^60^ and by extension the Rosetta Interface Score, does not fully include the entropic contribution to the global binding energy, which for peptides of this size tends to be substantial.^61^ Moreover, glycosylation has an effect on the dynamics of the peptide,^62^ in particular on the shape of the glycosylated peptide and the solvation/desolvation cost for the glucose moiety to unbind/bind to the protein, which is currently not considered.

### MD refinement

The best peptide-enzyme binding poses from the last stage of the docking protocol were used as starting conformations for MD simulations. The MD simulations were carried out in YASARA (www.yasara.org) in the AMBER14 force field.^63^ The protein-peptide complex was placed in a simulation box 10 Å larger than any complex atoms. TIP3P waters (∼28,000 molecules), ions (Na^+^ / Cl^-^, 0.15 M), and enough counterions (Na^+^) to neutralize the system were added to the simulation cell. The simulation was carried out using a multiple time step algorithm with a simulation time step of 1.25 fs.^64^ Electrostatics were handled with the PME algorithm with a cutoff of 7.86 Å.^65^ Pressure was kept at 1.0 bar with a Berendsen barostat,^66^ and temperature was maintained at 298 K with a modified Berendsen thermostat.^67^ The 10 ns simulations yielded a final fine-tuned peptide-enzyme complex.

### Data availability

The authors declare that data supporting the findings of this study are available within the paper and its supplementary information file. If not included in the supplementary information, the raw data (MS and MS/MS spectra, docking results and molecular simulations) will be made available by the corresponding author upon request.

## Supporting information

Supp Info Walvoort_NGT manuscript

## Acknowledgments

We thank the Interfaculty Mass Spectrometry Center of the University of Groningen (Dr. H.P. Permentier and Mr. M.P. de Vries) for help with whole-protein mass analysis, J. Hekelaar and M. Rovetta for their assistance with the site-preference experiments, Dr. R.H. Cool for help with the SPR analyses. We thank the Center for Information Technology of the University of Groningen for providing access to the Peregrine high-performance computing cluster. We thank Prof. B. Imperiali and Prof. G. Maglia for insightful discussions and proofreading of the manuscript. This work was financially supported by the Dutch Organization for Scientific Research (VENI 722.016.006) and the European Union through the Rosalind Franklin Fellowship COFUND project 60021 (both to M.T.C.W.), and by a CONACYT doctoral fellowship (to C.R.P.).

## Author contributions

L.Y. and M.T.C.W. conceived the project. L.Y., S.J.M. and M.T.C.W. designed and coordinated the study. The biochemical experiments were performed by L.Y., and the docking and simulations were performed by C.R.P. L.Y., C.R.P. and M.T.C.W. wrote the manuscript, with input from all authors. All authors were involved in proofreading the manuscript.

