## Supplementary material for "Semi-processive hyperglycosylation of adhesin by bacterial protein *N*-glycosyltransferases": Supp Info Walvoort_NGT manuscript

**Figure S1. Top:** Sequence of the C-terminal part of the HMW1 adhesin protein (HMW1ct). Asparagines in putative glycosylation sites (numbered) are in bold and underlined. **Bottom:** I-TASSER model of HMW1ct with glycosylation sites highlighted in yellow. Left: side view. Right: top view.

MAHHHHHHVWTANSGALTTLAGSTIKGTESVTTSSQSGDIGGTISGGTVEVKATES
LTTQSNSKIKATTGEA**<u>N</u>(1)**VTSATGTIGGTISGNTV**<u>N</u>(2)**VTANAGDLTVGNGAEI**<u>N</u>(3)**
ATEGAATLTTSSGKLTTEASSHITSAKGQV**<u>N</u>(4)**LSAQDGSVAGSINAA**<u>N</u>(5)**VTL**<u>N</u>(6)**
TTGTLTTVKGSNI**<u>N</u>(7)**ATSGTLVINAKDAELNGAALG**<u>N</u>(8)**HTVV**<u>N</u>(9)**ATNA**<u>N</u>(10)**GS
GSVIATTSSRV**<u>N</u>(11)**ITGDLITINGLNIISKNGINTVLLKGVKIDVKYIQPGIASVDEVIEA
KRILEKVKDLSDEEREALAKLGVSAVRFIEP**<u>N</u>(12)**NTITVDTQNEFATRPLSRIVISEG
RACFSNSDGATVCVNIADNGR

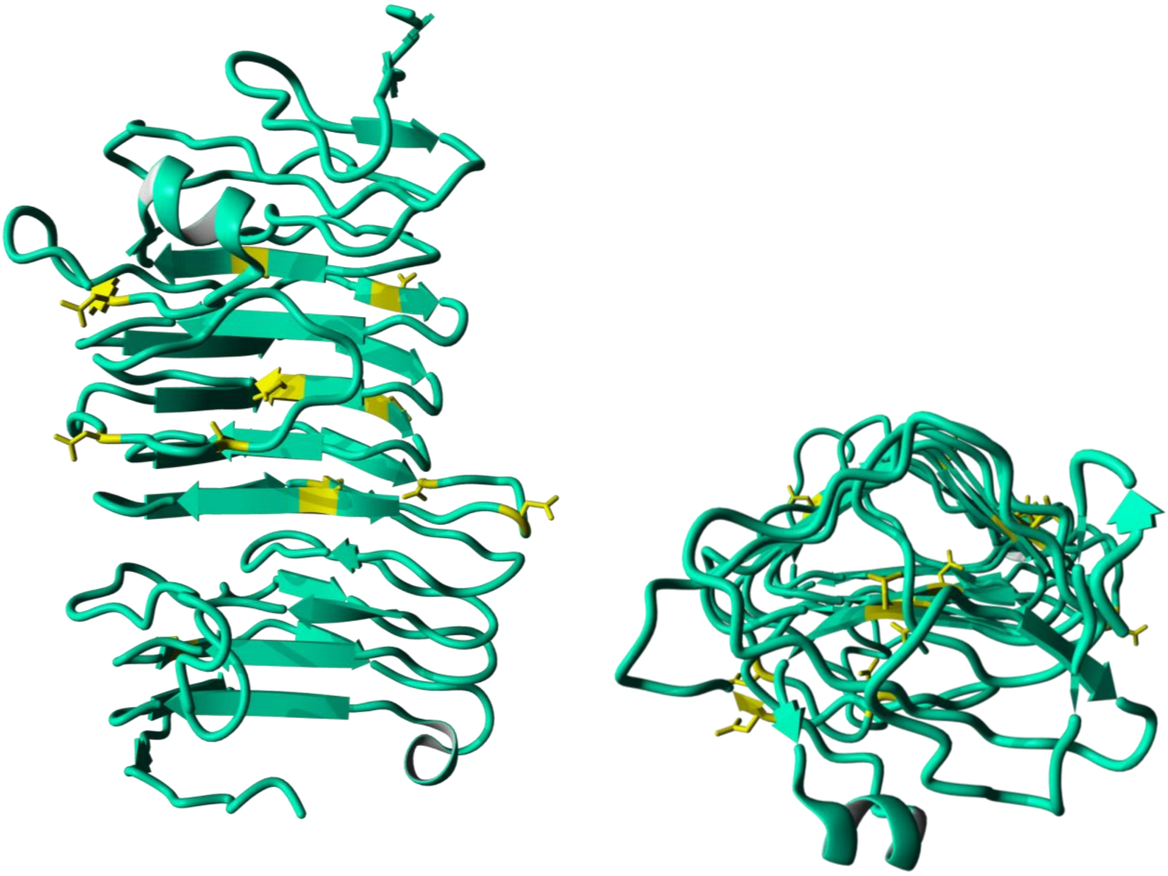

**Figure S2.** Product profiles from the time-course of ApNGT-HMW1ct glycosylation at 1:10 ratio after 1 min (A) and 5 min (B) and 15 h (C).

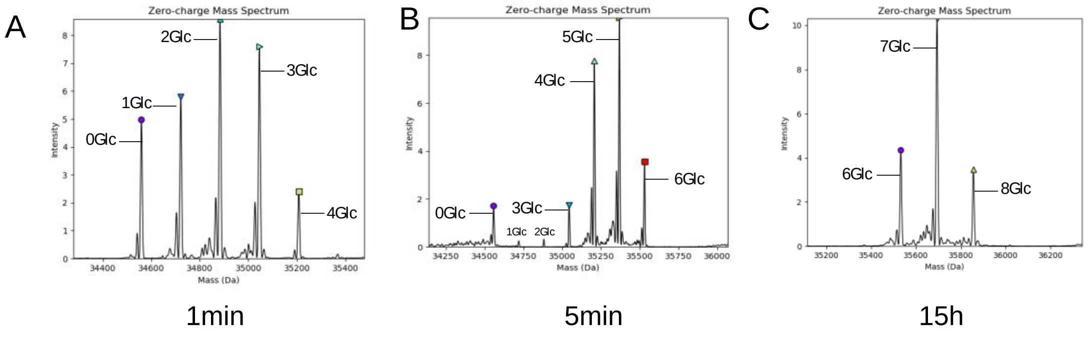

**Figure S3.** Product profiles from the time-course of ApNGT-HMW1ct glycosylation at 1:100 ratio after 10 min (A) and 15 h (B).

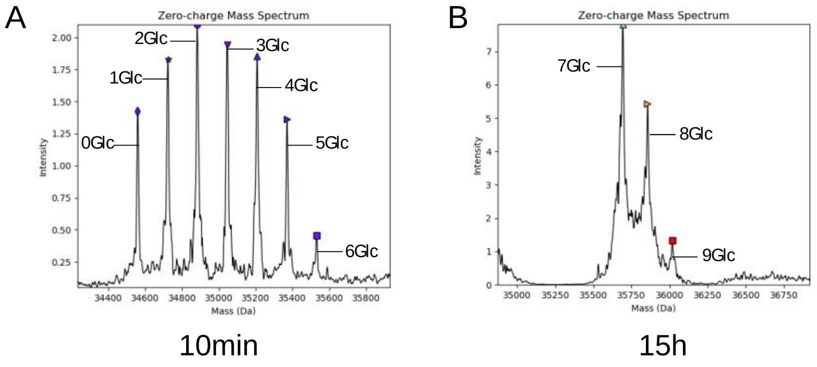

**Figure S4.** Product profile in the timecourse of HiNGT-HMW1ct glycosylation at 1:10 ratio after 5 min (A) 10 min (B), 30 min (C), 90 min(D), 300 min (E) and 15 h (F).

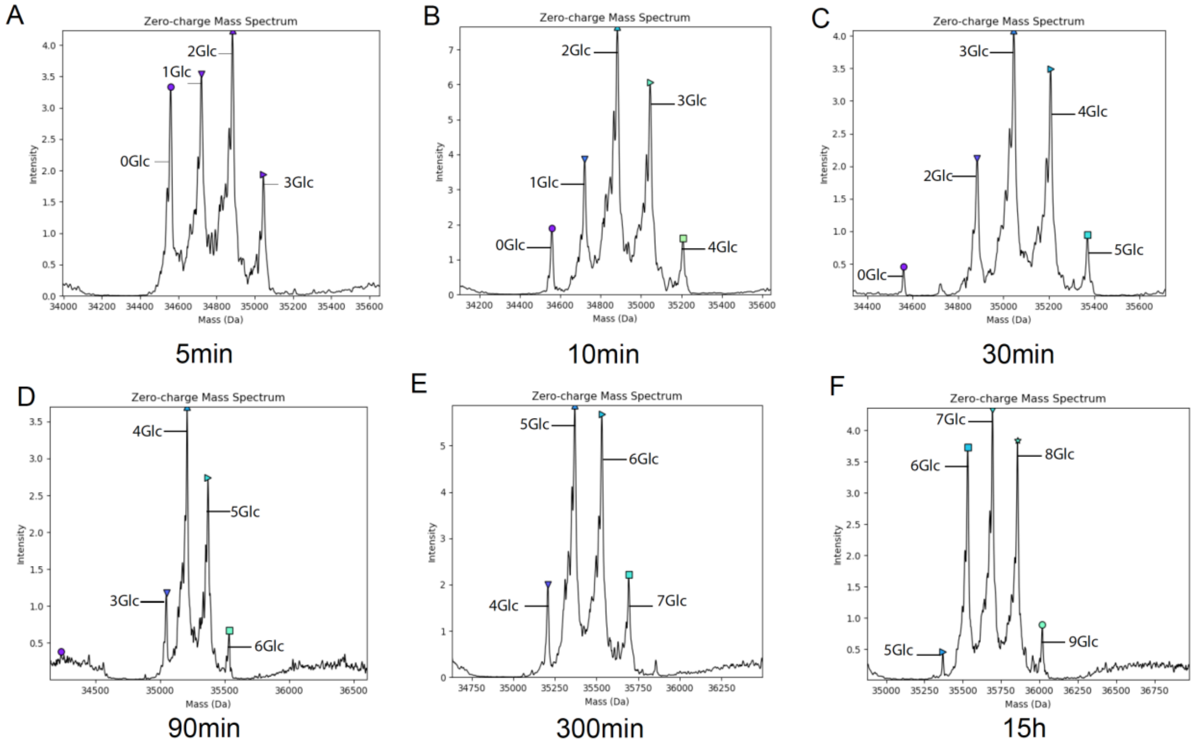

**Figure S5.** Product profile in the timecourse of HiNGT-HMW1ct glycosylation at 1:100 ratio after 180 min.

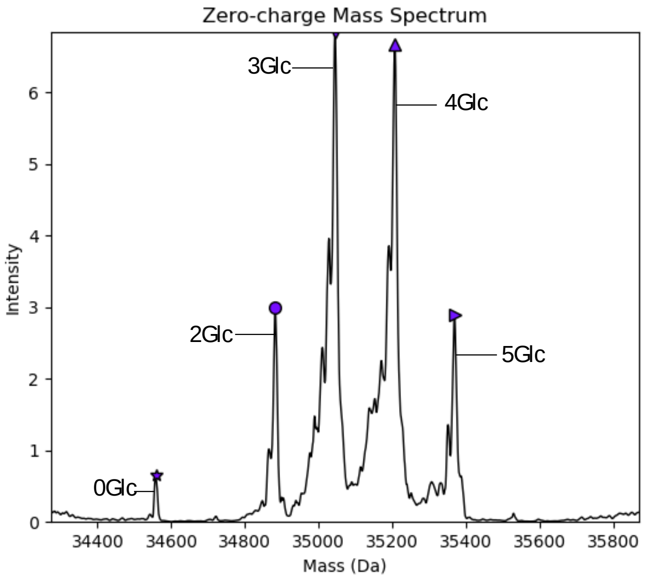

**Table S1** Processivity parameter calculations for ApNGT-HMW1ct reaction, 1:100 ratio, at 10min.

| Number of Glc added | % active NGT at n Glc | $P_n$ |
|---|---|---|
| **1** | 100 | 0.,22 |
| **2** | 22.01 | 0.92 |
| **3** | 20.34 | 0.95 |
| **4** | 19.39 | 0.74 |
| **5** | 14.26 | 0.34 |
| **6** | 4.82 | |

**Table S2** Processivity parameter calculations for HiNGT-HMW1ct reaction, 1:100 ratio, after 30min.

| Number of Glc added | % active NGT at n Glc | $P_n$ |
|---|---|---|
| **1** | 100 | 0.42 |
| **2** | 41.95 | 0.59 |
| **3** | 24.85 | 0.1 |
| **4** | 2.49 | |

**Figure S6. Non-linear regression fit.** $k_{cat}$ and $K_m$ were determined for ApNGT by performing the continuous coupled-assay with increasing concentration of ApHMW1ct, and the initial velocities were fitted using Graphpad Prism. Initial velocities were averaged over n=8, and concentration ApNGT was constant at 100 nM.

1

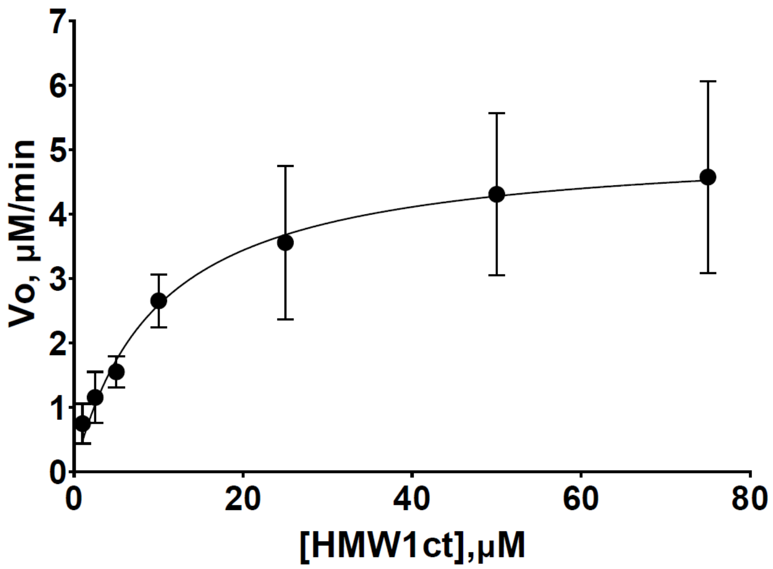

| Nonlin fit<br>Table of results | | A<br>Vo, uM/min |
|---|---|---|
| 1 | kcat | |
| 2 | Best-fit values | |
| 3 | Et | = 0.1000 |
| 4 | kcat | 51.12 |
| 5 | Km | 9.722 |
| 6 | Vmax | = 5.112 |
| 7 | 95% CI (profile likelihood) | |
| 8 | kcat | 44.68 to 59.11 |
| 9 | Km | 6.092 to 15.64 |
| 10 | Goodness of Fit | |
| 11 | Degrees of Freedom | 54 |
| 12 | R squared | 0.7347 |
| 13 | Sum of Squares | 41.06 |
| 14 | Sy.x | 0.8720 |
| 15 | Constraints | |
| 16 | Et | Et = 0.1 |
| 17 | | |
| 18 | Number of points | |
| 19 | # of X values | 56 |
| 20 | # Y values analyzed | 56 |

**Figure S7. Kinetic analysis of HiNGT.** Continuous coupled-assay with increasing concentration of ApHMW1ct, and the initial velocities were fitted using Graphpad Prism. Initial velocities were averaged over n=4, and concentration HiNGT was constant at 100 nM.

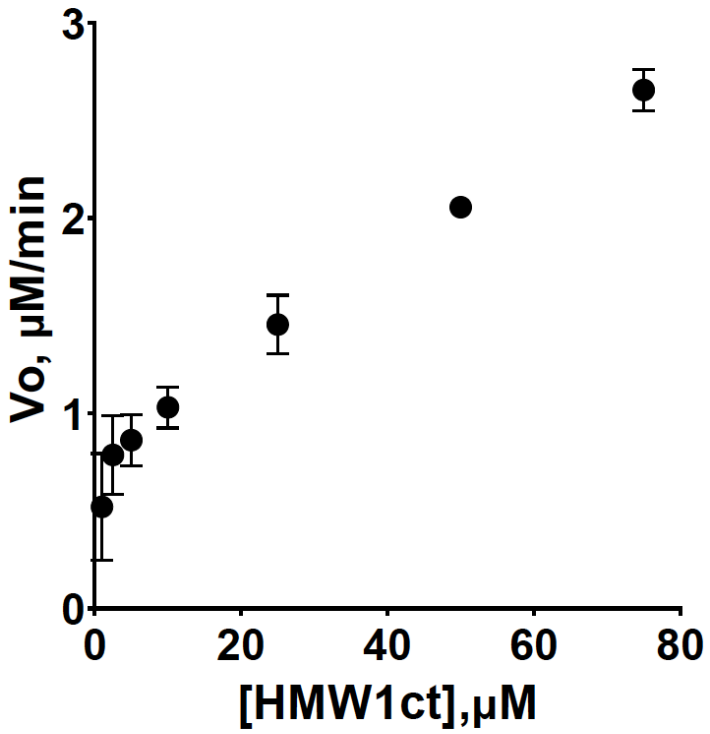

**Figure S8.** SPR sensograms for association-dissociation events between ApNGT and HMW1ct (**A**) and Glc-HMW1ct (**B**)

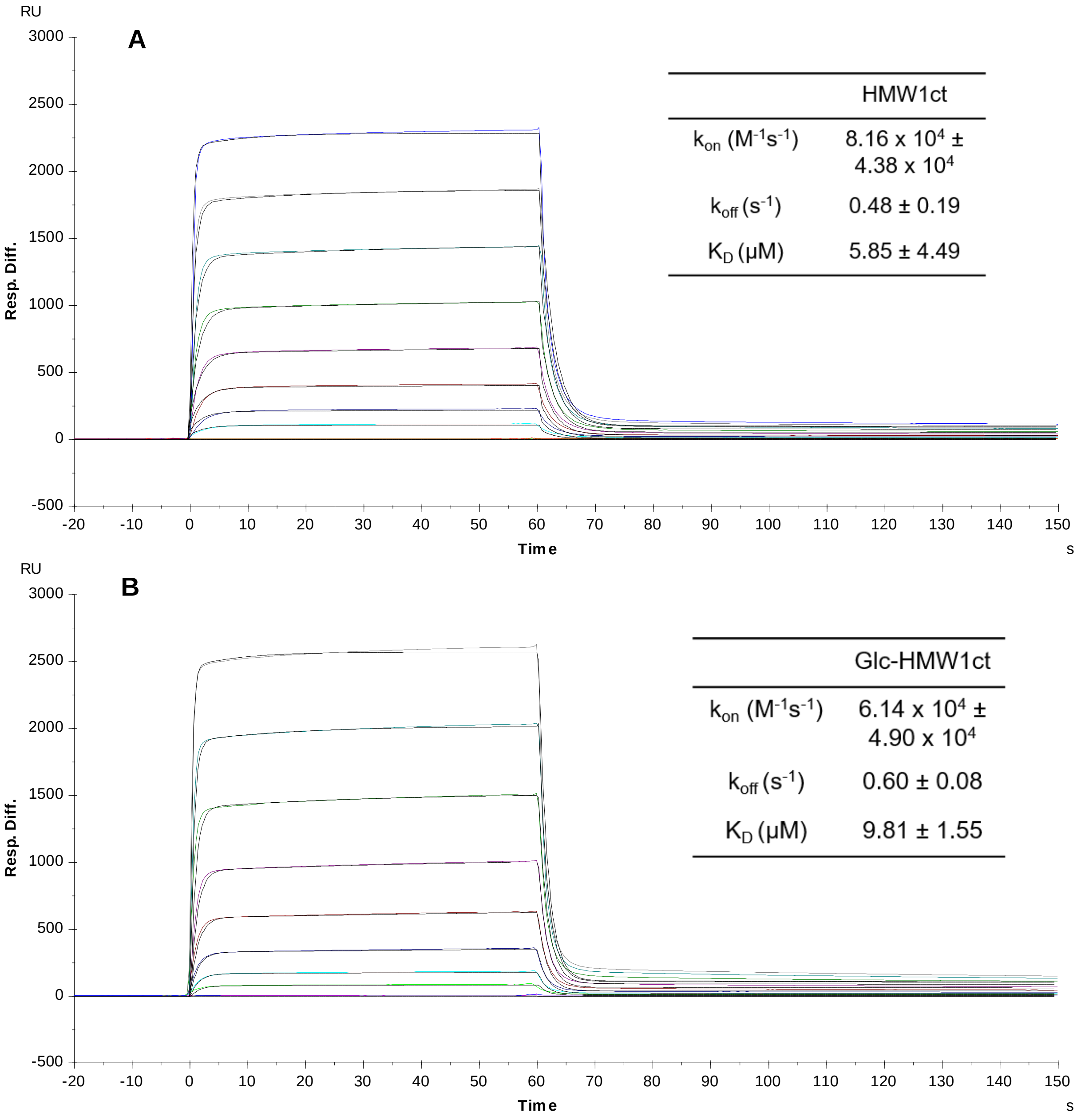

| | HMW1ct |
|---|---|
| $k_{on}$ ($M^{-1}s^{-1}$) | $8.16 \times 10^4 \pm 4.38 \times 10^4$ |
| $k_{off}$ ($s^{-1}$) | $0.48 \pm 0.19$ |
| $K_D$ (μM) | $5.85 \pm 4.49$ |

| | Glc-HMW1ct |
|---|---|
| $k_{on}$ ($M^{-1}s^{-1}$) | $6.14 \times 10^4 \pm 4.90 \times 10^4$ |
| $k_{off}$ ($s^{-1}$) | $0.60 \pm 0.08$ |
| $K_D$ (μM) | $9.81 \pm 1.55$ |

**Figure S9. Effect of different enzyme-substrate concentrations within the same ratio on glycosylation product profile.** For ApNGT-catalyzed glycosylation the ratio 1:100 of enzyme to substrate was used (A), and for the ApNGT-catalyzed glycosylation the ratio 1:10 of enzyme to substrate was used (B). A fixed concentration of **1 mM UDP-Glc** was used. Data presented are average of two independent experiments.

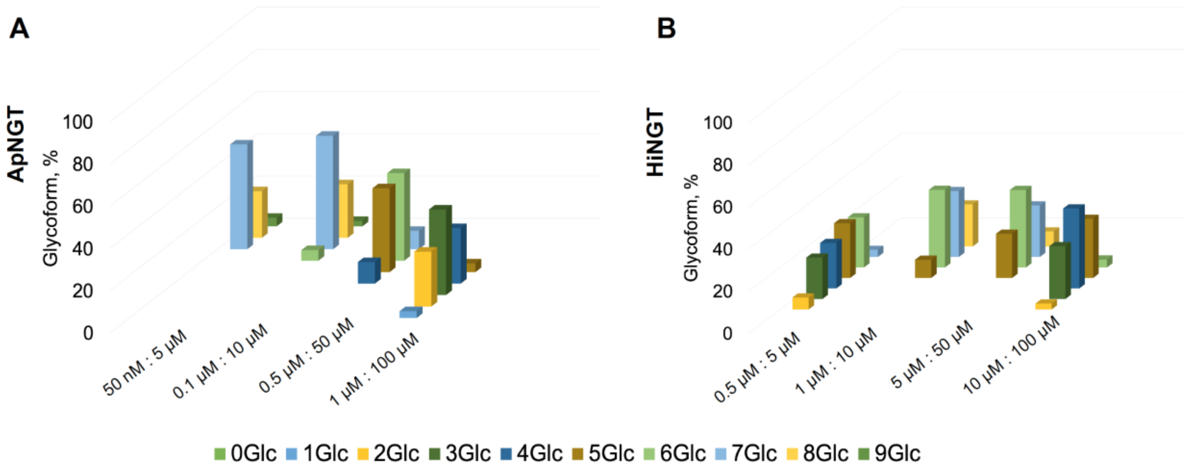

**Figure S10. Effect of different enzyme-substrate concentrations within the same ratio on glycosylation product profile.** For ApNGT-catalyzed glycosylation the ratio 1:100 of enzyme to substrate was used (A), and for the ApNGT-catalyzed glycosylation the ratio 1:10 of enzyme to substrate was used (B). The amount of **UDP-Glc was used in excess** to the substrate, resulting in 1 mM for 5 μM and 10 μM HMW1ct, 5 mM for 50 μM HMW1ct, and 10 mM for 100 μM HMW1ct. Data presented are average of two independent experiments.

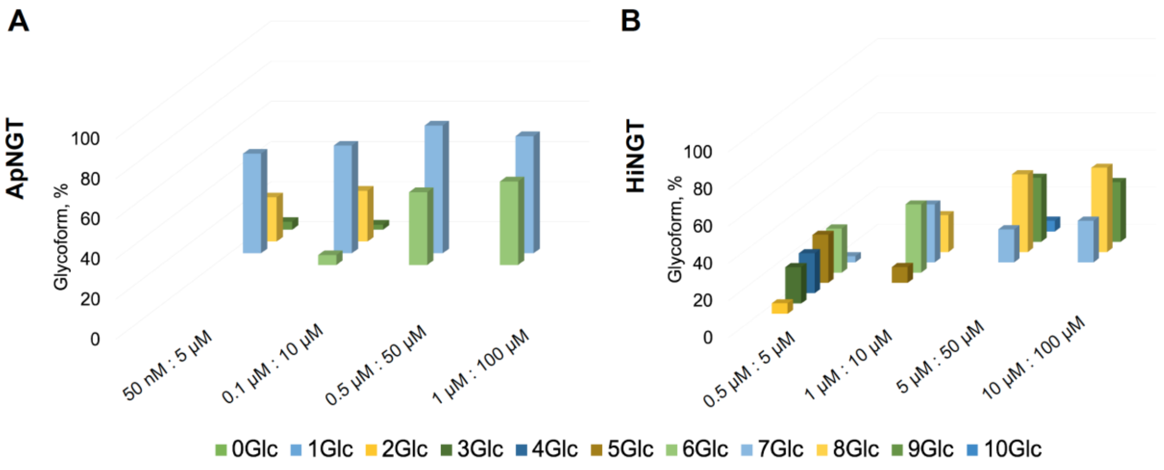

**Figure S11. Distraction assay.** Product profile of ApNGT-HMW1ct glycosylation after addition of product after 10min (A). Product profile of HiNGT-HMW1ct glycosylation after addition of product after 10min (B).

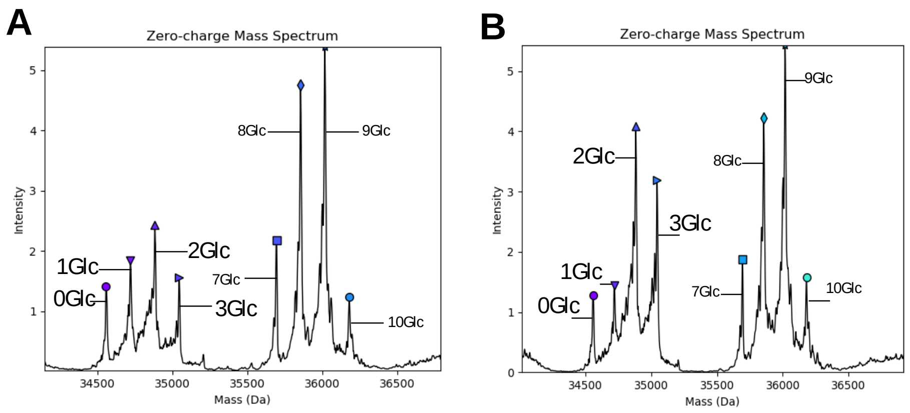

**Figure S12. Restarting the overnight reaction by addition of substrate (*top*) or early glycoforms (*bottom*). A, C:** ApNGT-catalyzed reaction. **B, D:** HiNGT-catalyzed reaction. The overnight reaction (10 μM HMW1ct, 1:50 ratio for ApNGT and 1:5 ratio for HiNGT) and an equal volume of 20 μM of HMW1ct or 20 μM of early glycoforms (EG, separately generated) was added to reach the desired ratios (1:100 for ApNGT and 1:10 for HiNGT). Conclusions are based on data of two independent experiments. Deconvolved mass spectra are shown below.

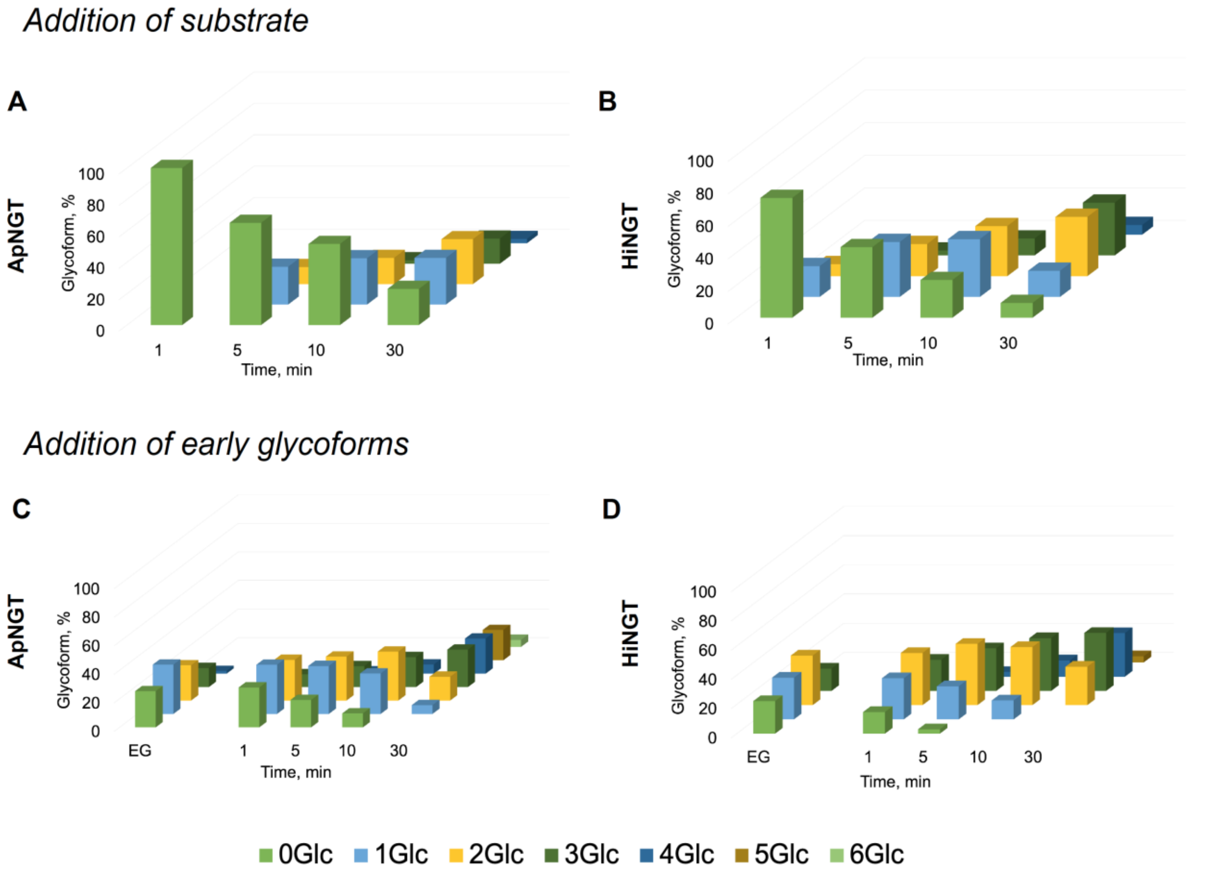

A (left): early glycoforms generated in ApNGT-HMW1ct reaction. A (right): early glycoforms generated in HiNGT-HMW1ct reaction. Product profiles of the restarted ApNGT-HMW1ct reaction after 30min of addition of early glycoforms (B) or substrate (C). Product profiles of the restarted HiNGT-HMW1ct reaction after 30min of addition of early glycoforms (D) or substrate (E).

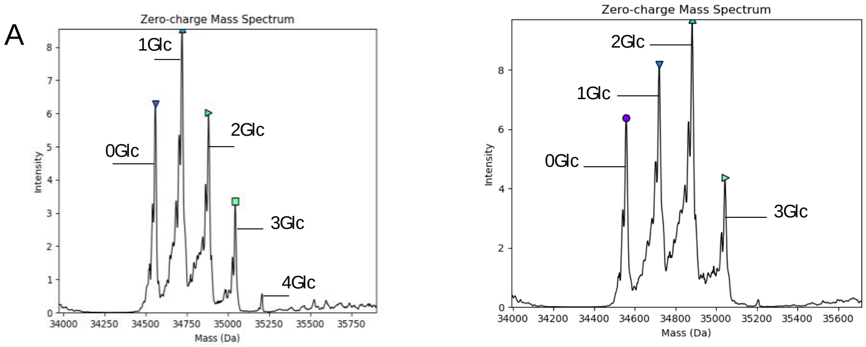

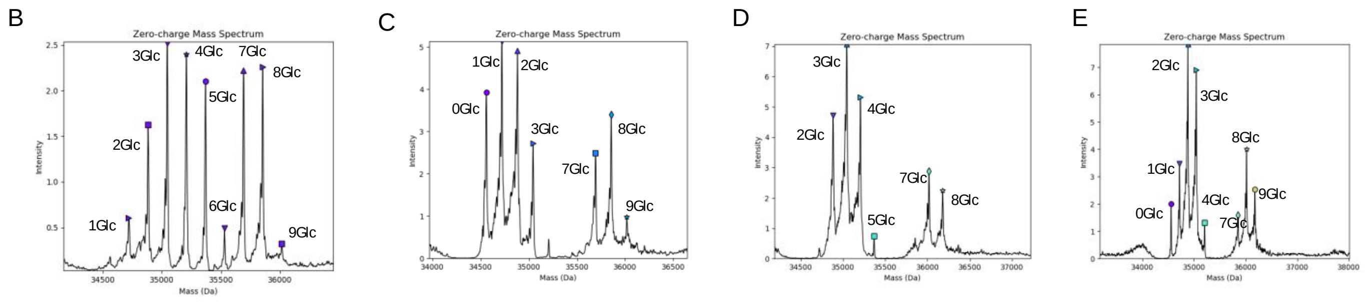

Restarting ApHMW1C-HMW1ct reaction with early glycoforms 30min

Restarting ApHMW1C-HMW1ct reaction with substrate 30min

Restarting HiHMW1C-HMW1ct reaction with early glycoforms 30min

Restarting HIHMW1C-HMW1ct reaction with substrate 30min

**Figure S13.** Product profile of the full timecourse of ApNGT (A) and HiNGT (B) catalyzed HMW1ct glycosylation under the single-hit conditions (1:500 ratio).

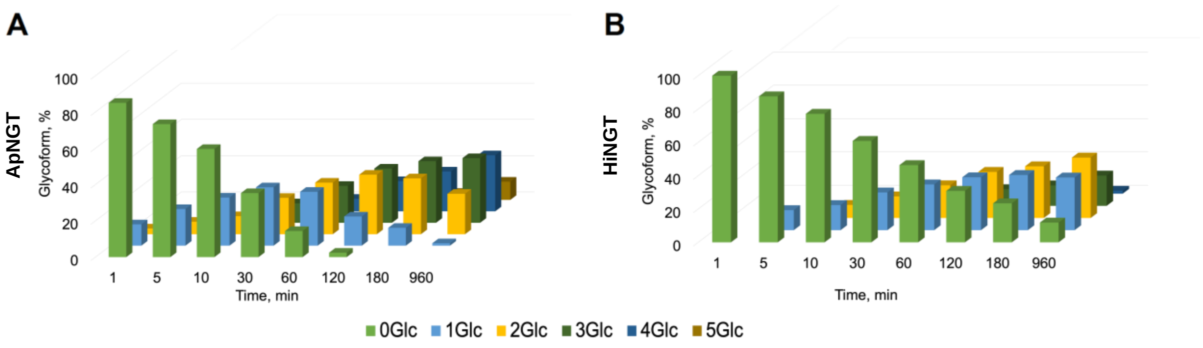

**Figure S14.** Product profile of the full timecourse of ApNGT (A) and HiNGT (B) catalyzed HMW1ct glycosylation under the single-hit conditions (1:1000 ratio).

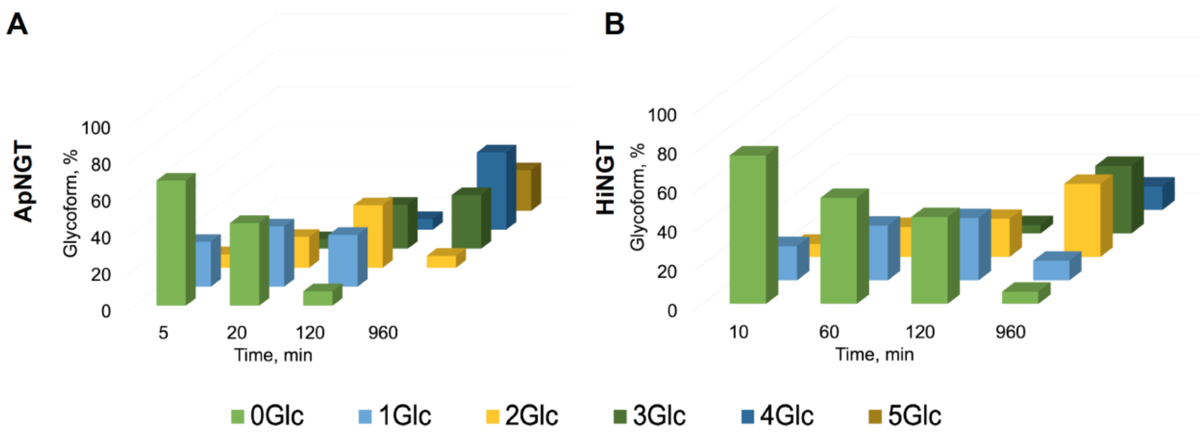

**Figure S15. Preference for N-glycosylation sites in HMW1ct, later time points.**

A) Site-specific modification for ApNGT after 2.5 min; B) Site-specific modification for HiNGT after 20 min; C) I-TASSER model of HMW1ct with sequon sites (yellow) and non-sequon sites (magenta).

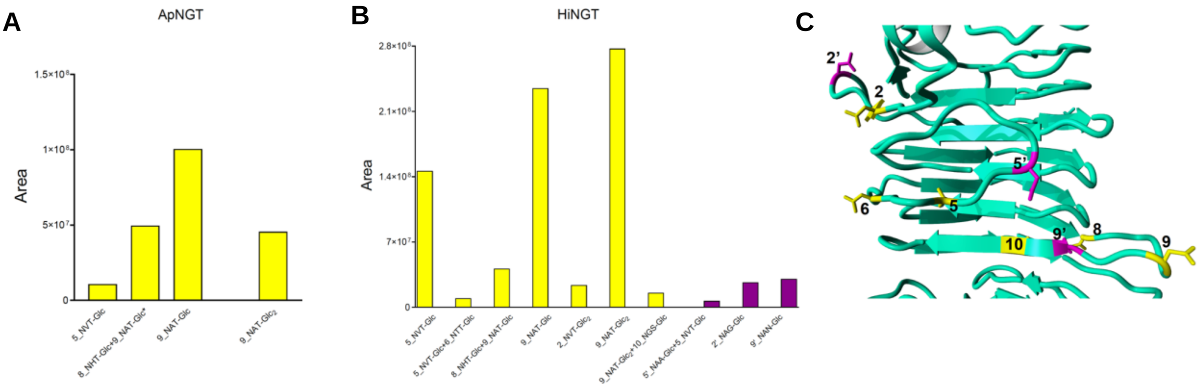

**Figure S16. Peptide spectra for ApNGT site preference at the 0.5 min time point.**

1. 8_NHT-Glc+9_NAT-Glc

GSNINATSGTLVINAKDAELNGAALG**N**HTVV**N**ATNANGSGSVIATTSSR

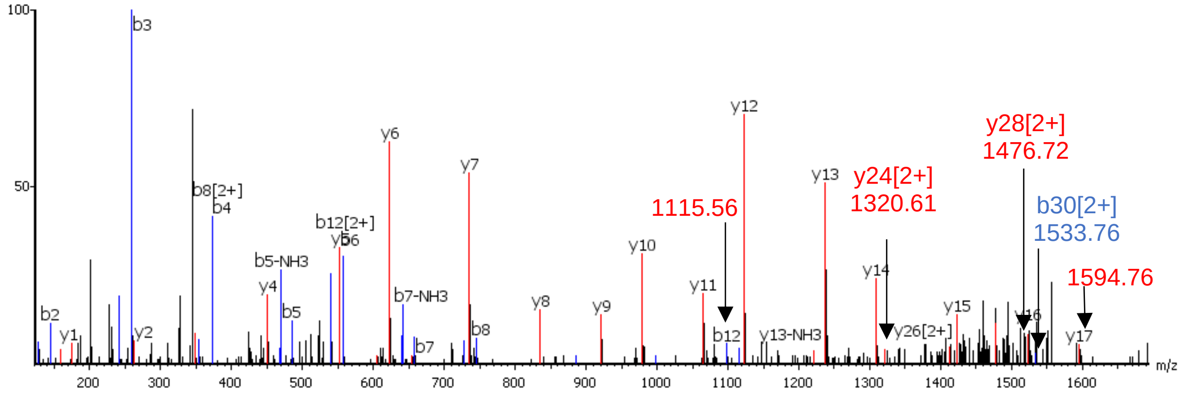

2. 9_NAT-Glc

DAELNGAALGNHTVV**N**ATNANGSGSVIATTSSR

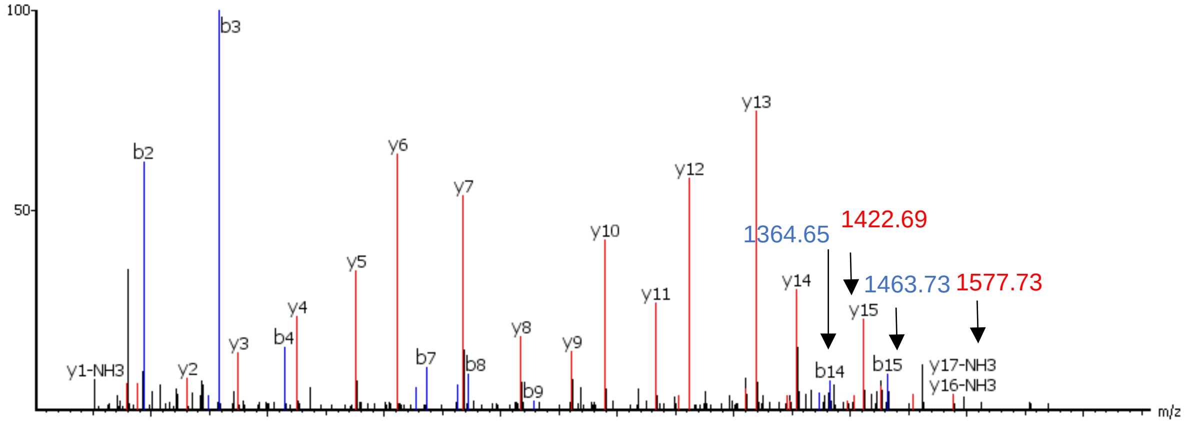

3. 5'_NAA-Glc

GQVNLSAQDGSVAGSI**N**AANVTLNTTGTLTTVK

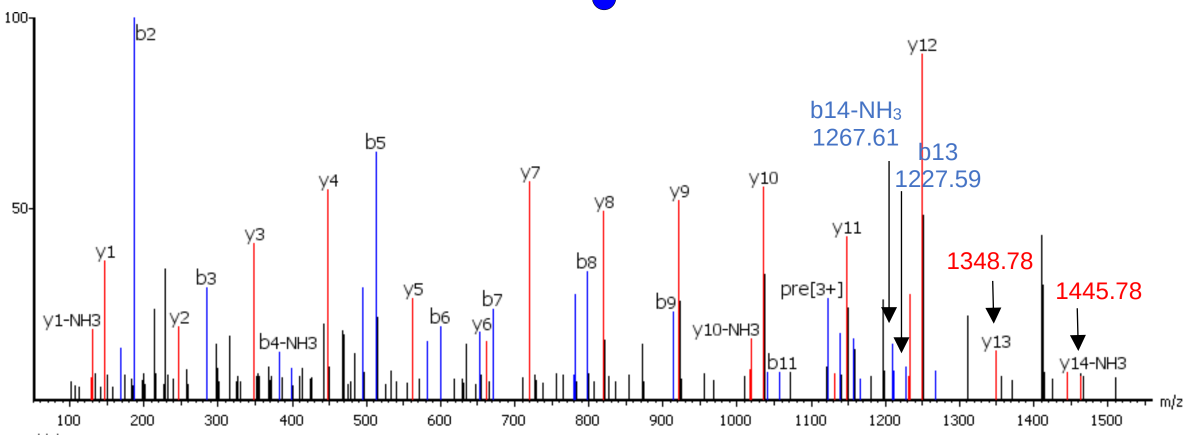

**Figure S17. Peptide spectra for ApNGT site preference at the 2.5 min time point.**

1. 5_NVT-Glc

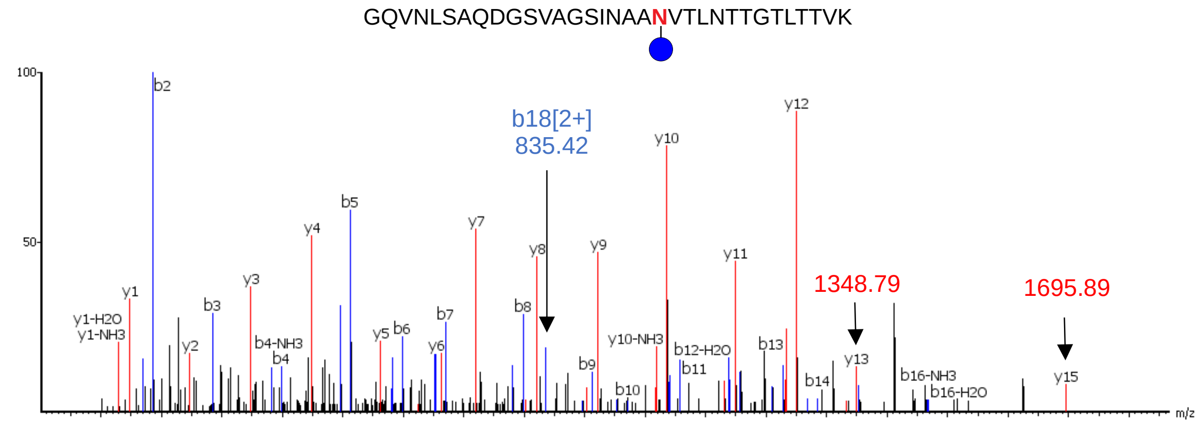

2. 8_NHT-Glc+9_NAT-Glc

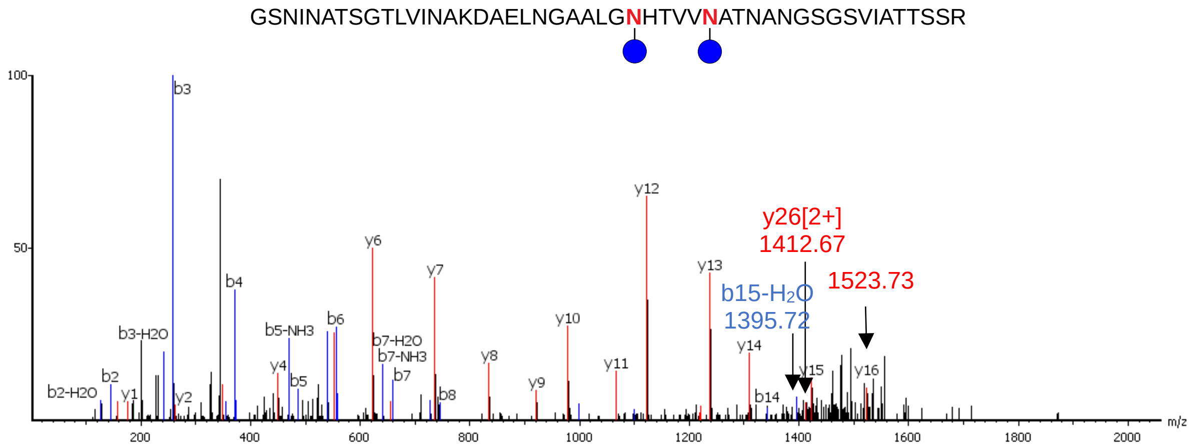

3. 9_NAT-Glc

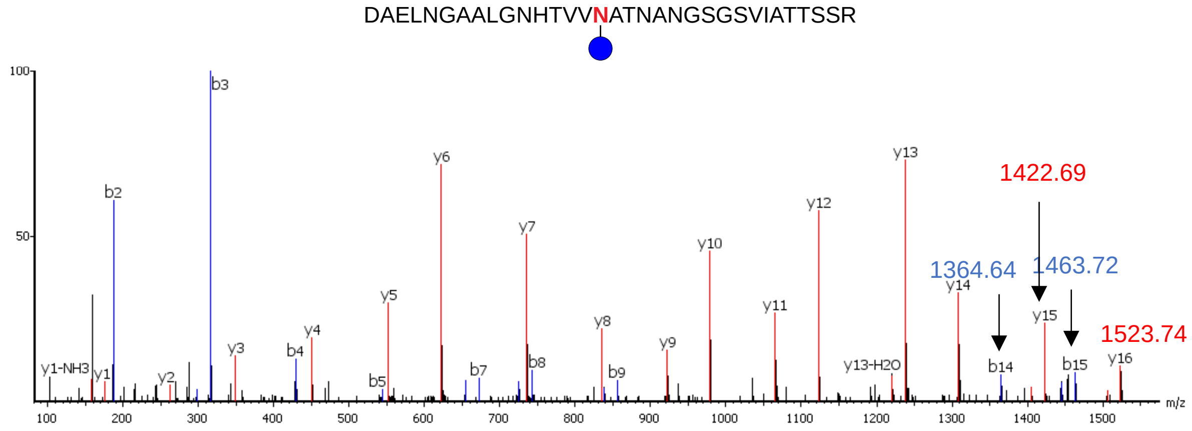

4. 9_NAT-$Glc_2$

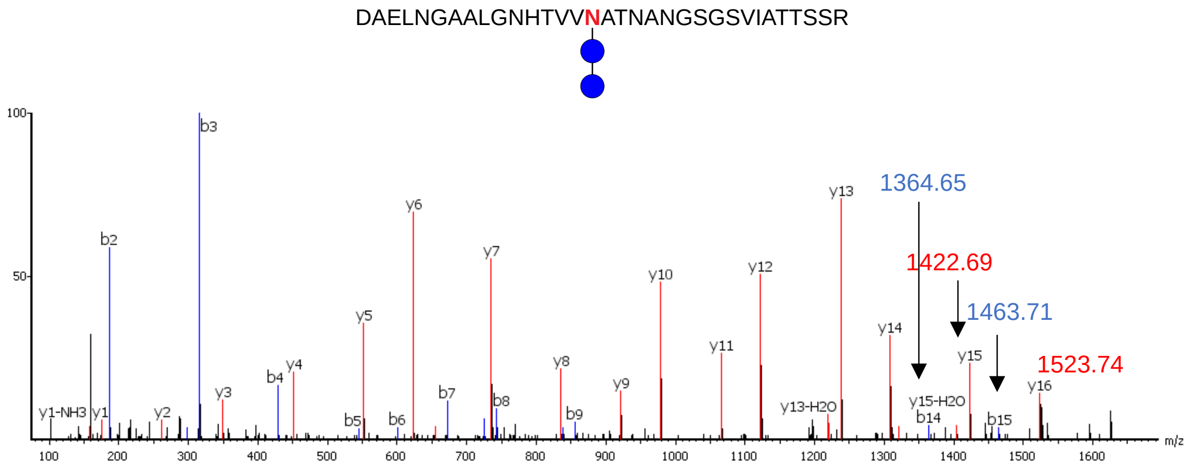
DAELNGAALGNHTVVNATNANGSGSVIATTSSR
1364.65
1422.69
1463.71
1523.74
y1-NH3
y1
b2
y2
b3
y3
b4
y4
b5
y5
b6
y6
b7
y7
b8
y8
b9
y9
y10
y11
y12
y13
y13-H2O
y14
y15-H2O
b14
y15
b15
y16
m/z

**Figure S18. Peptide spectra for HiNGT site preference at the 0.5 min time point.**

1. 9_NAT-Glc

DAELNGAALGNHTVV**N**ATNANGSGSVIATTSSR

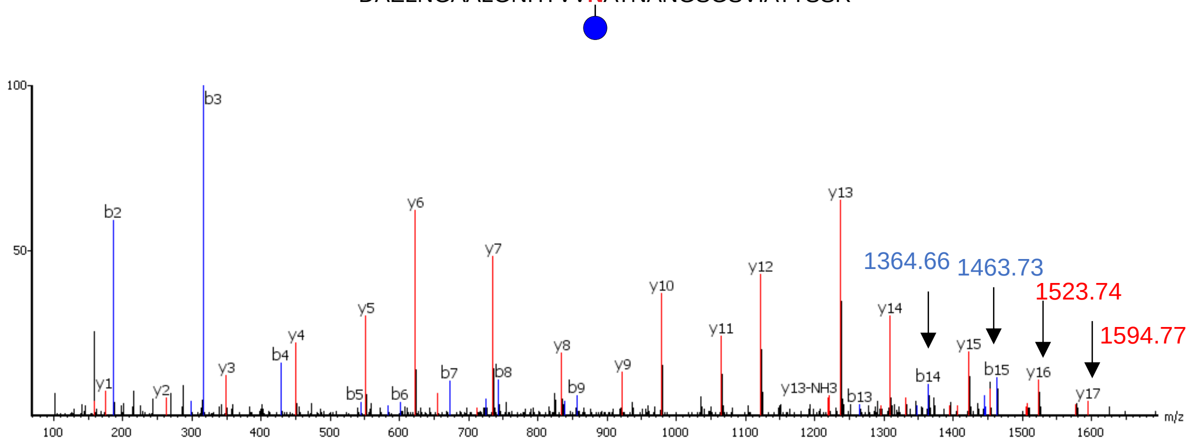

2. 2'_NAG

ATTGEANVTSATGTIGGTISGNTVNVTA**N**AGDLTVGNGAEINATEGAATLTTSSGK

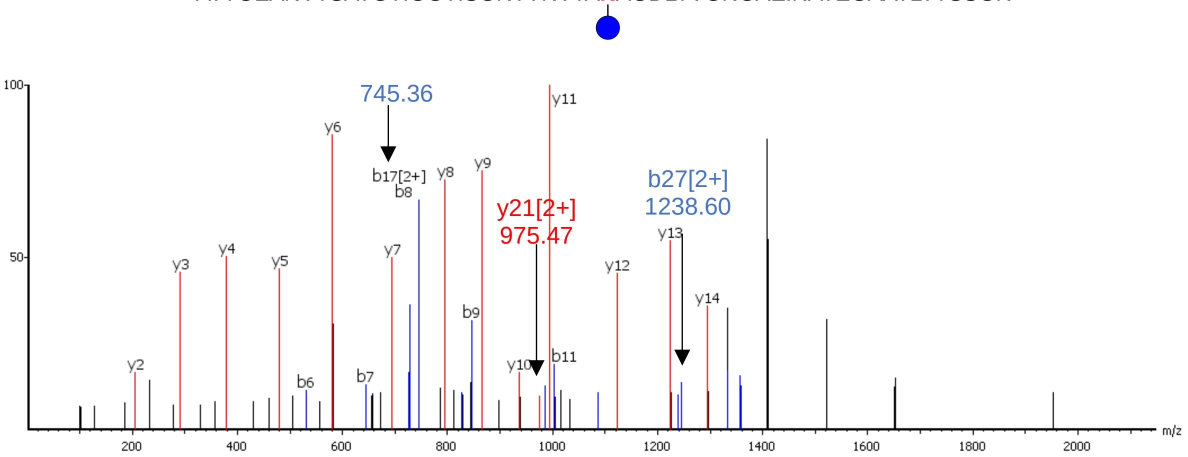

3. 9_NAT-$Glc_2$

DAELNGAALGNHTVV**N**ATNANGSGSVIATTSSR

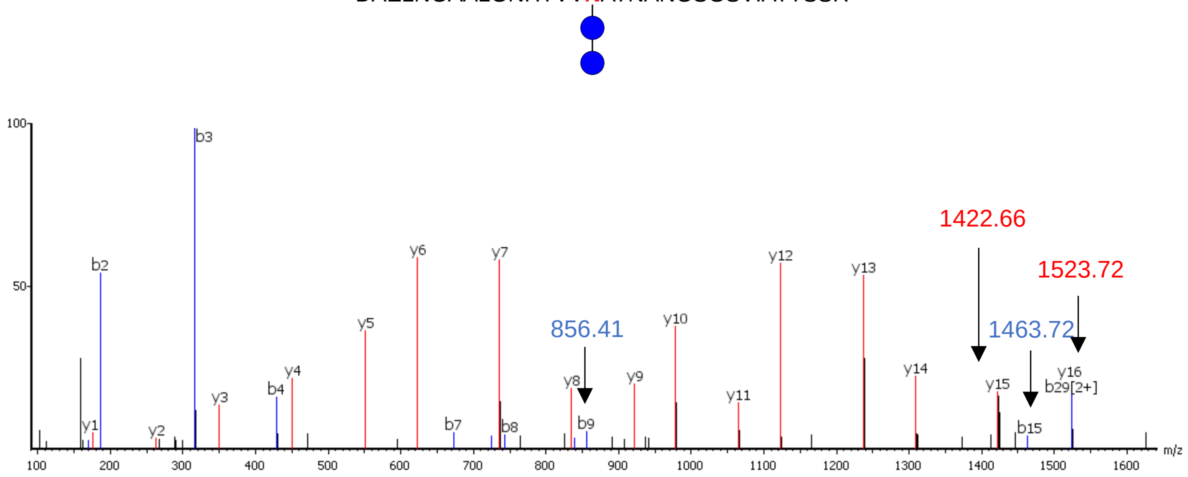

4. 9'_NAN-$Glc_2$

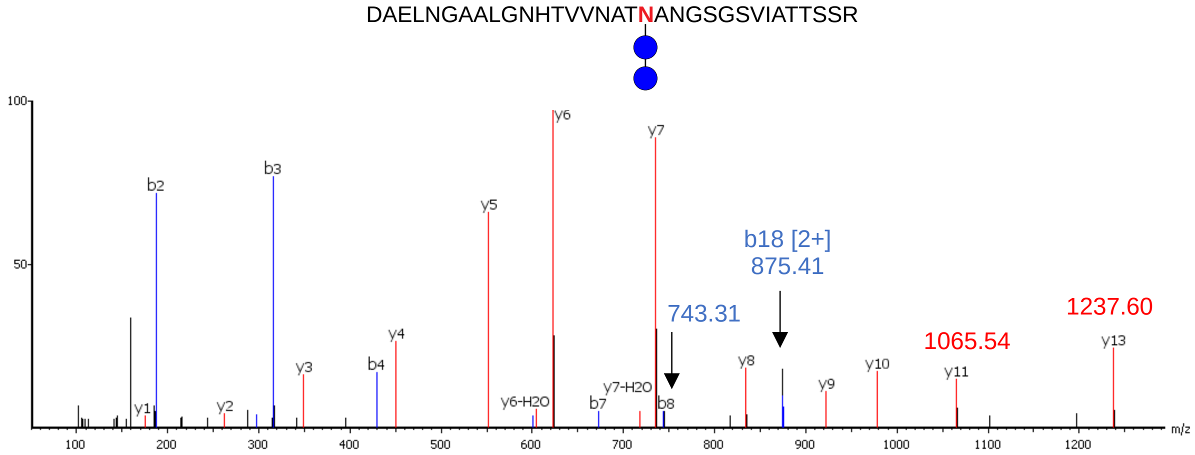

DAELNGAALGNHTVVNATNANGSGSVIATTSSR
b18 [2+]
875.41
743.31
1065.54
1237.60
b2
b3
b4
b7
b8
y1
y2
y3
y4
y5
y6
y7
y8
y9
y10
y11
y13
y6-H2O
y7-H2O
m/z

**Figure S19. Peptide spectra for HiNGT site preference at the 20 min time point.**

1. 5_NVT-Glc

GQVNLSAQDGSVAGSINAA**N**VTLNTTGTLTTVK

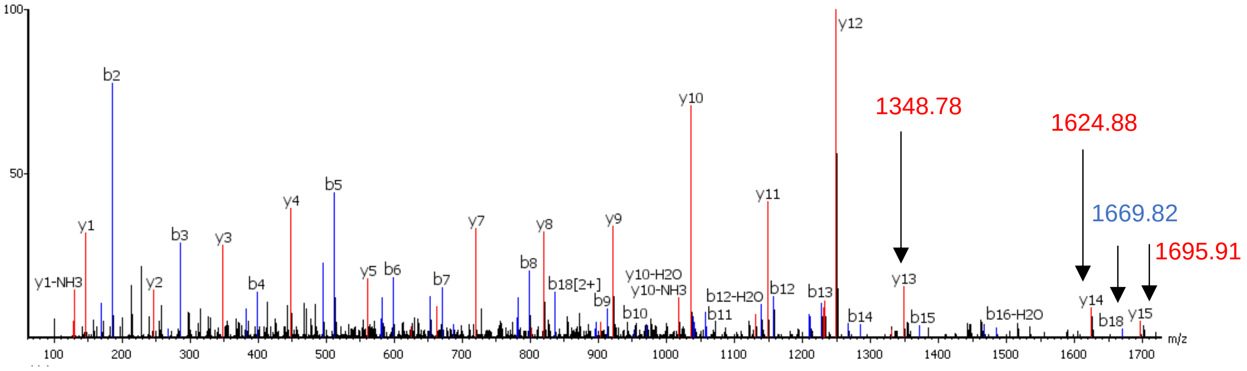

2. 5_NVT-Glc + 6_NTT-Glc

GQVNLSAQDGSVAGSINAA**N**VTL**N**TTGTLTTVK

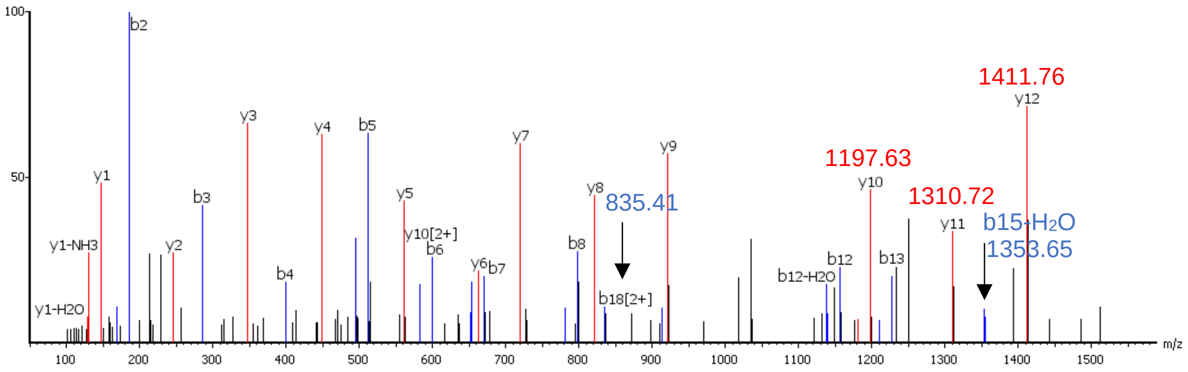

3. 8_NHT-Glc + 9_NAT-Glc

GSNINATSGTLVINAKDAELNGAALG**N**HTVV**N**ATNANGSGSVIATTSSR

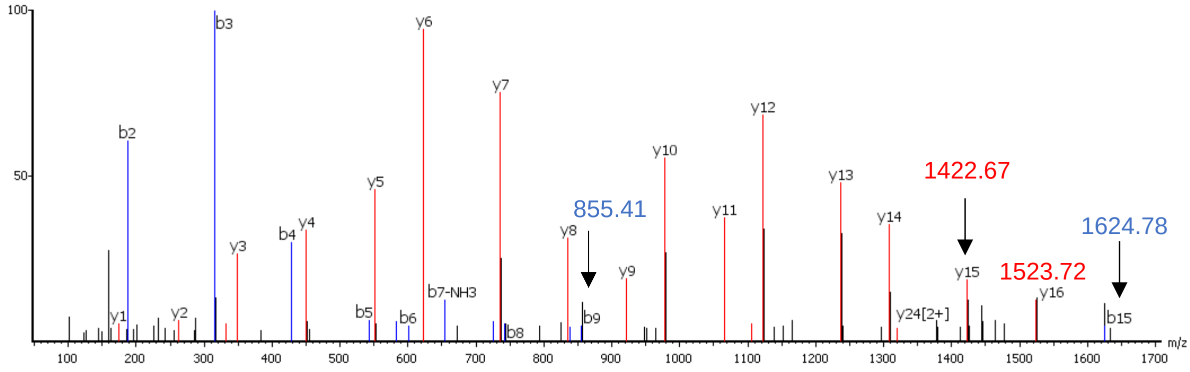

4. 9_NAT-Glc

DAELNGAALGNHTVVNATNANGSGSVIATTSSR

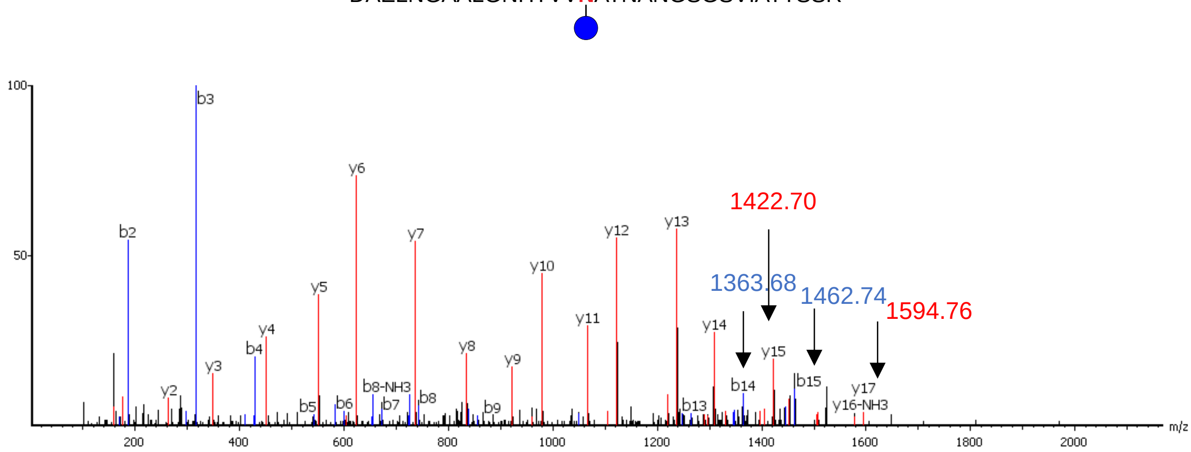

5. 2_NVT-Glc2

ATTGEANVTSATGTIGGTISGNTVNVTANAGDLTVGNGAEINATEGAATLTTSSGK

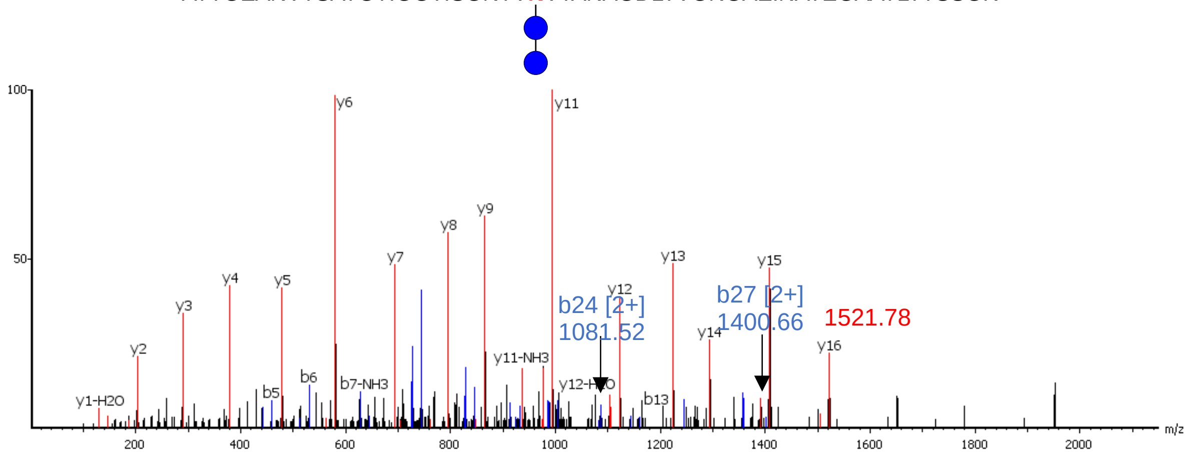

6. 9_NAT-Glc2

DAELNGAALGNHTVVNATNANGSGSVIATTSSR

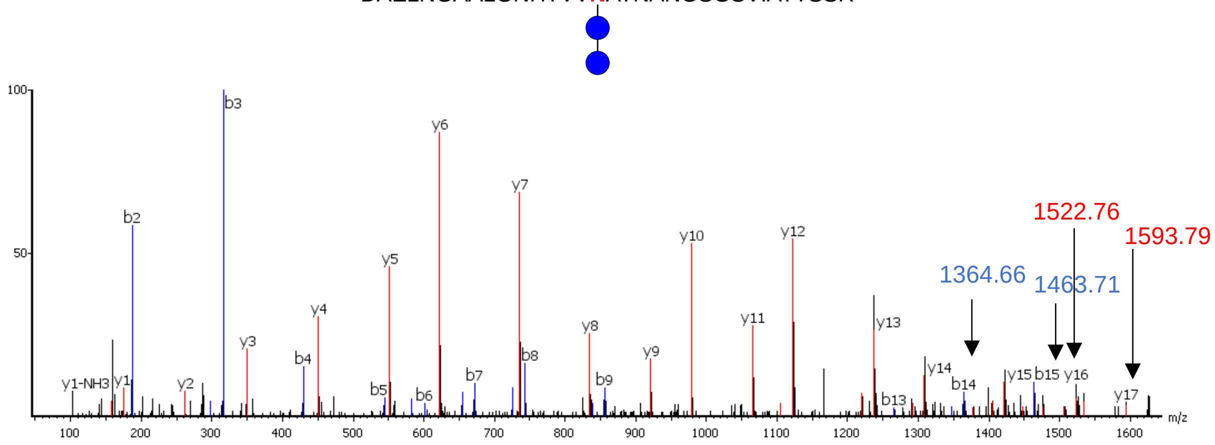

7. 9_NAT-Glc$_2$ + 10_NGS-Glc

DAELNGAALGNHTVVNATNANGSGSVIATTSSR

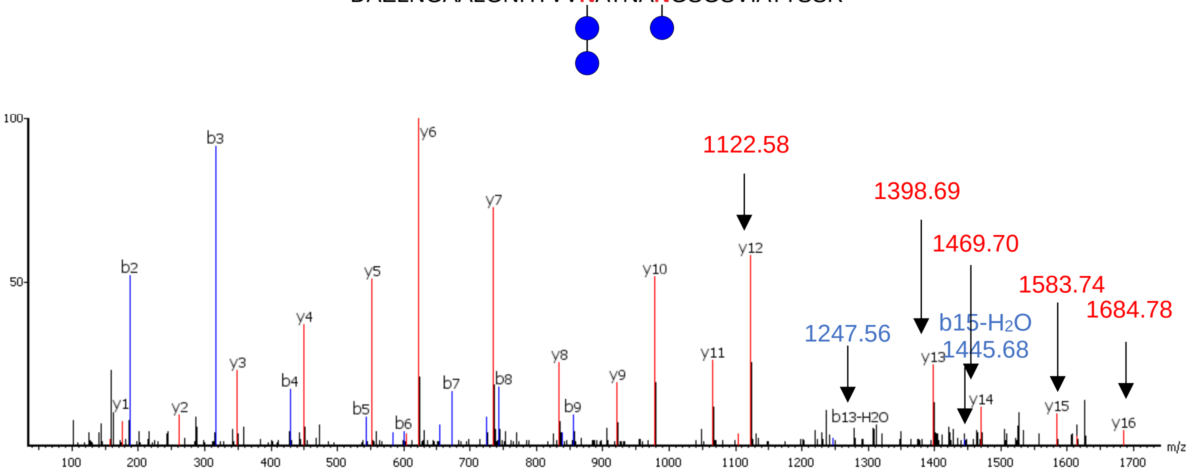

8. 5'_NAA-Glc + 5_NVT

GQVNLSAQDGSVAGSINAANVTLNTTGTLTTVK

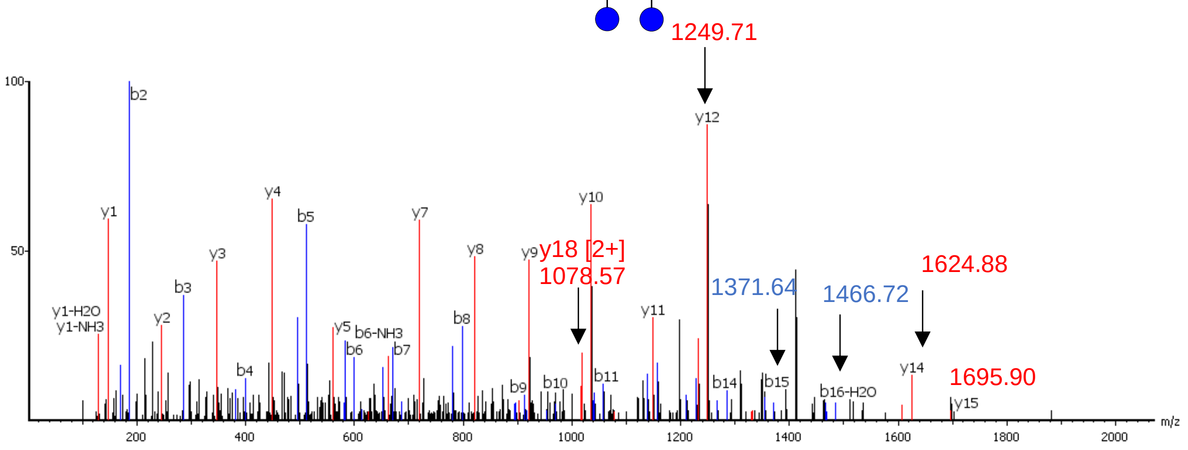

9. 2'_NAG-Glc

ATTGEANVTSATGTIGGTISGNTVNVTANAGDLTVGNGAEINATEGAATLTTSSGK

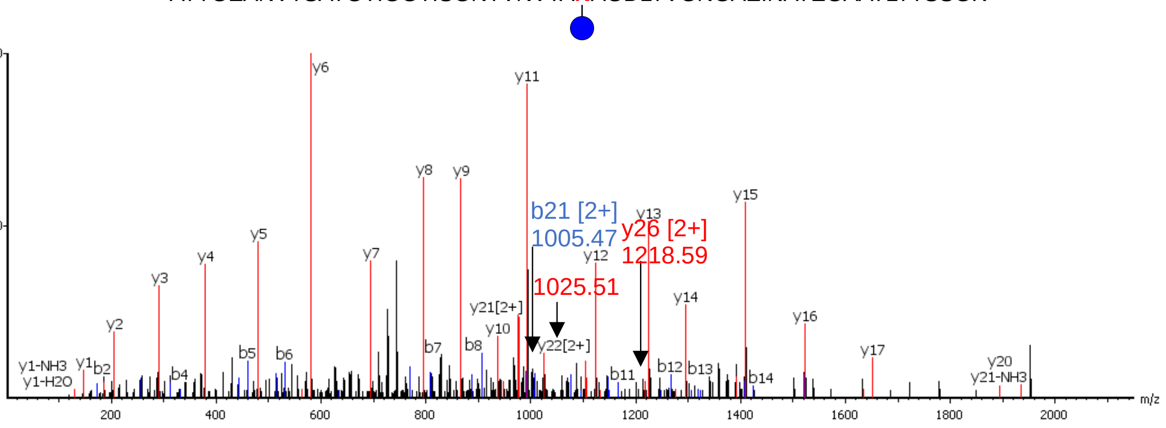

10. 9’_NAN-Glc

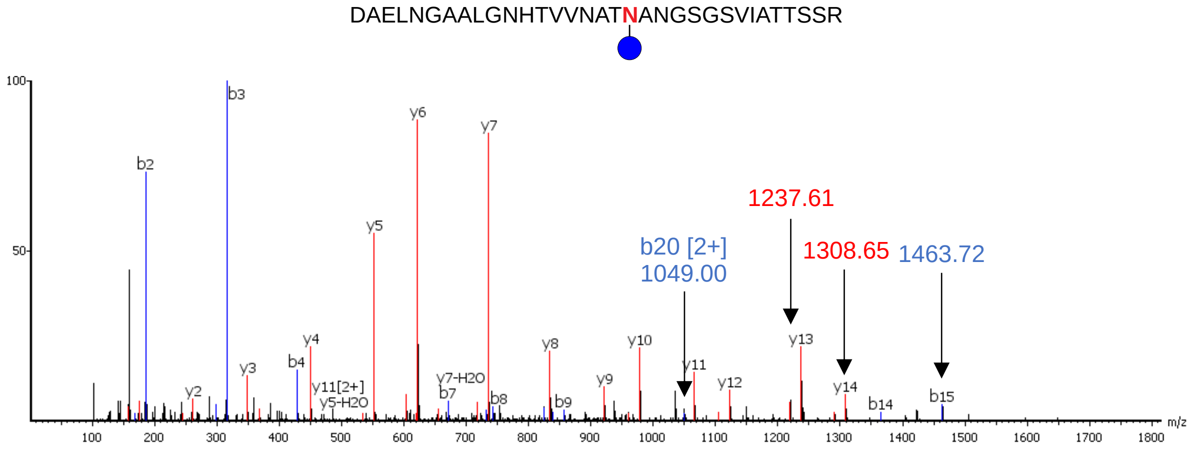

DAELNGAALGNHTVVNATNANGSGSVIATTSSR
b3
y6
y7
b2
y5
1237.61
b20 [2+]
1049.00
1308.65
1463.72
y4
y8
y10
y13
b4
y3
y11
y12
y14
y2
y11[2+]
y5-H2O
y7-H2O
b7
b8
b9
y9
b14
b15
m/z

**Figure S20. Docking of the ApNGT::UDP-Glc complex**. Structure of the ApNGT::UDP-Glc complex obtained by computational modelling shows important protein residues around the Glc moiety, and a comparison to the crystal structure of hOGT::UDP-GlcNAc (PDB code: 4GZ5).

The structure of ApNGT in complex with UDP-Glc places the sugar in a cavity formed by residues Tyr222, Ile279, Gly370, His371, and Lys441, while the anomeric position ($C_\alpha$) of UDP-Glc remains accessible to nucleophilic attack from the solvent phase. Comparing the nucleotide-sugar binding pose to the hOGT::UDP-GlcNAc complex, as the only other member of the GT41 enzyme family (PDB: 4GZ5) shows that the nucleotide-sugars adopt a similar conformation with the sugar moieties placed in a similar protein environment (Tyr222/Tyr841, His371/His498, Gly370/Gly654, Lys441/Lys842, Gln469/Gln839).

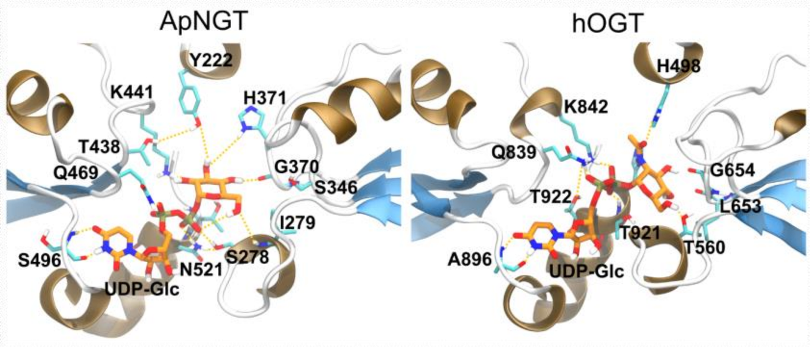

**Figure S21. hOGT::peptide::UDP-GlcNAc complex.** Two binding modes for hOGT from crystal structures. The green cartoon corresponds to PDB codes 6MA3, 6MA2, 6MA5, 6MA4, 4N3A, and 4N39. The purple cartoon corresponds to PDB codes 5HGV, 3PE4, 4GYW, 4GZ3, 4N3B, and 4N3C.

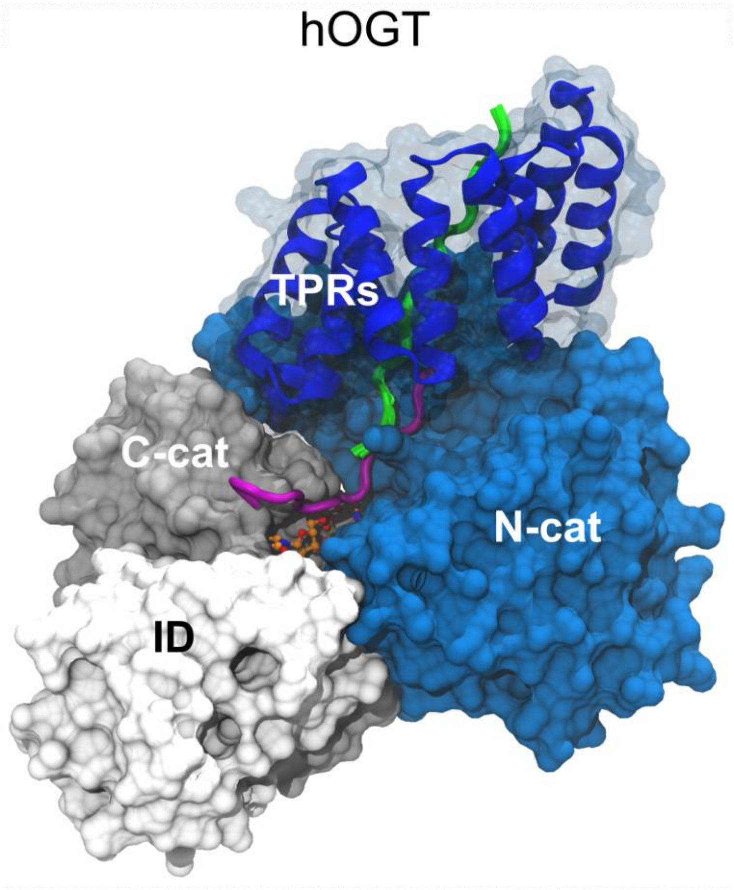

**Figure S22. Sequence alignment of ApNGT and HiNGT** (created using CLC Sequence Viewer version 7)

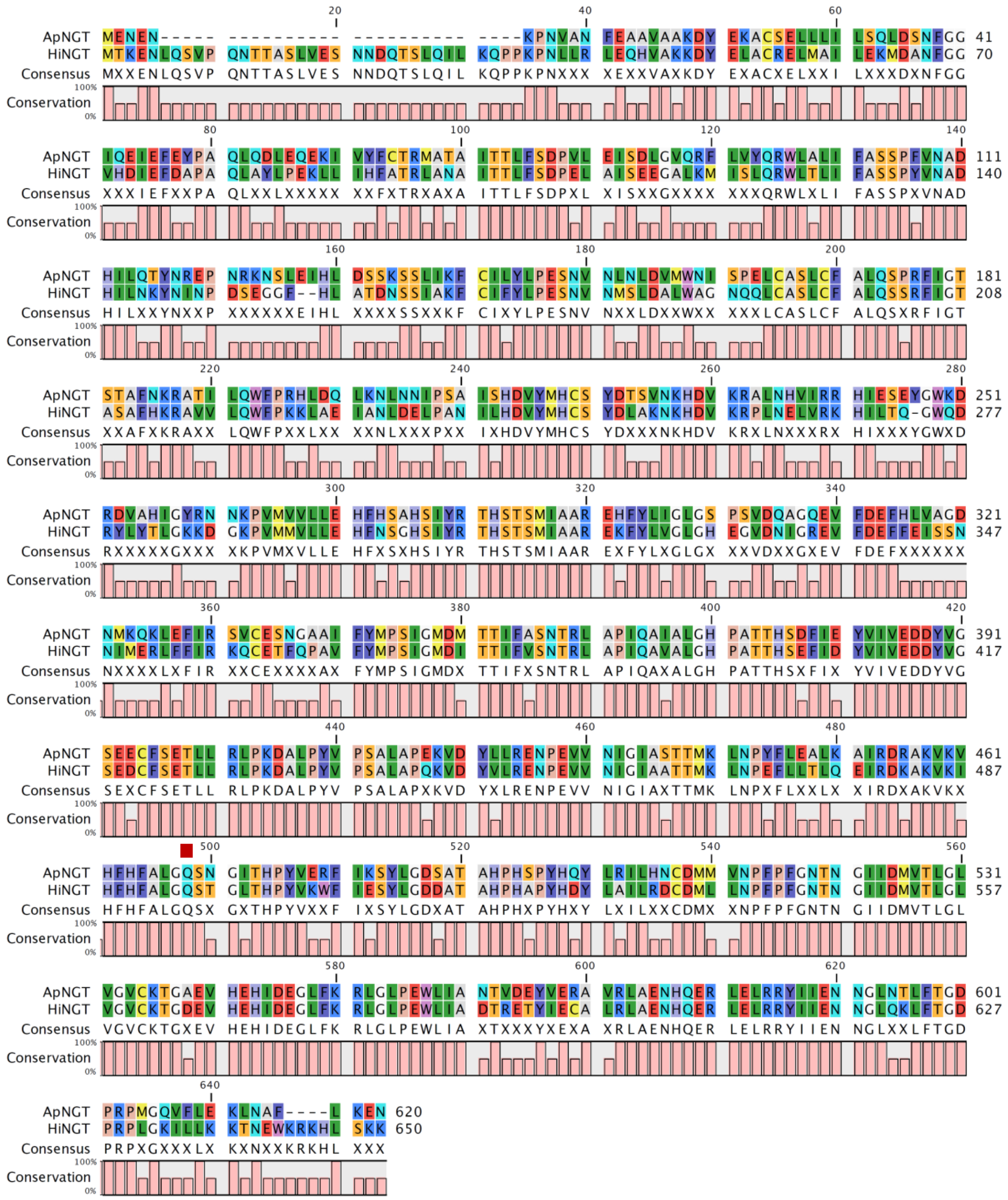

**Table S3: Settings used in the deconvolution using UniDec.**

| | |
|---|---|
| **Data processing:** | |
| m/z range | 600 to 3000 |
| Subtract minimum | 0.0 |
| Gaussian smoothing | 0.0 |
| Bin every | 0.1 |
| **Additional Data Processing Parameters**<br>**Unidec parameters:** | |
| Charge Range | 20 to 100 |
| Mass range | 10000 to 100000 |
| Sample Mass Every (Da) | 1 |
| Peak FWHM (Th) | 0.1 |
| Peak Shape Function | Gaussian |
| **Additional Filters/Restraints**<br>**Peak Selection and Plotting:** | |
| Peak Detection Range (Da) | 50.0 |
| Peak Detection Threshold | 0.03 |
| Peak Normalization | None |

**Table S4.** Calculated and experimentally found masses of HMW1ct glycoforms.

| Glycoform | Calculated mass | Experimental mass |
|---|---|---|
| 0 | 34560 | 34559 |
| 1 | 34722 | 34721 |
| 2 | 34884 | 34882 |
| 3 | 35046 | 35044 |
| 4 | 35208 | 35207 |
| 5 | 35370 | 35369 |
| 6 | 35532 | 35530 |
| 7 | 35694 | 35695 |
| 8 | 35856 | 35856 |
| 9 | 36018 | 36017 |
| 10 | 36180 | 36180 |

**Figure S23. Comparison of ionization intensities** of 10 µM HMW1ct (**A**), 10 µM Glc-HMW1ct (**B**), and a mixture of 10 µM HMW1ct + 10 µM Glc-HMW1ct (**C**)

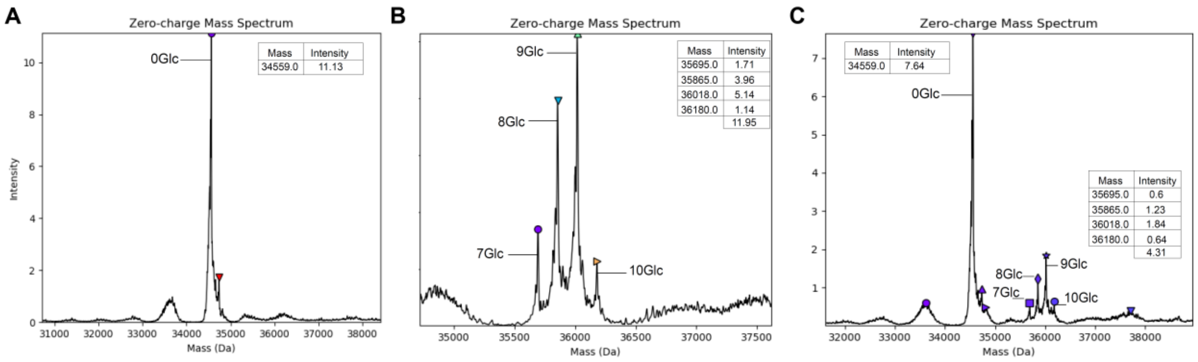

### Table S5. HMW1ct tryptic peptides identified in proteomics analysis.

| Sequon sites | Peptides |
|---|---|
| Site 1 | IKATTGEA**N(+162.05)V**TSATGTIGGTISGNTVNVTANAGDLTVGNGAEINATEGAATLTTSSGK |
| | ATTGEA**N(+162.05)VT**SATGTIGGTISGNTVNVTANAGDLTVGNGAEINATEGAATLTTSSGK |
| Site 2 | IKATTGEANVTSATGTIGGTISGNTV**N(+162.05)VT**ANAGDLTVGNGAEINATEGAATLTTSSGK |
| | ATTGEANVTSATGTIGGTISGNTV**N(+162.05)VT**ANAGDLTVGNGAEINATEGAATLTTSSGK |
| Site 3 | IKATTGEANVTSATGTIGGTISGNTVNVTANAGDLTVGNGAEI**N(+162.05)AT**EGAATLTTSSGK |
| | ATTGEANVTSATGTIGGTISGNTVNVTANAGDLTVGNGAEI**N(+162.05)AT**EGAATLTTSSGK |
| Site 4 | LTTEASSHITSAKGQV**N(+162.05)LS**AQDGSVAGSINAANVTLNTTGTLTTVK |
| | GQV**N(+162.05)LS**AQDGSVAGSINAANVTLNTTGTLTTVK |
| Site 5 | LTTEASSHITSAKGQVNLSAQDGSVAGSINAA**N(+162.05)VT**LNTTGTLTTVK |
| | GQVNLSAQDGSVAGSINAA**N(+162.05)VT**LNTTGTLTTVK |
| Site 6 | LTTEASSHITSAKGQVNLSAQDGSVAGSINAANVTL**N(+162.05)TT**GTLTTVK |
| | GQVNLSAQDGSVAGSINAANVTL**N(+162.05)TT**GTLTTVK |
| Site 7 | GSNI**N(+162.05)AT**SGTLVINAKDAELNGAALGNHTVVNATNANGSGSVIATTSSR |
| | GSNI**N(+162.05)AT**SGTLVINAK |
| Site 8 | GSNINATSGTLVINAKDAELNGAALG**N(+162.05)HT**VVNATNANGSGSVIATTSSR |
| | DAELNGAALG**N(+162.05)HT**VVNATNAN(+.98)GSGSVIATTSSR |
| Site 9 | GSNINATSGTLVINAKDAELNGAALGNHTVV**N(+162.05)AT**NANGSGSVIATTSSR |
| | DAELNGAALGNHTVV**N(+162.05)AT**NANGSGSVIATTSSR |
| Site 10 | DAELNGAALGNHTVVNATNA**N(+162.05)GS**GSVIATTSSR |

| Non-sequon sites | Peptides |
|---|---|
| Site 1' | ATTGEANVTSATGTIGGTISG**N(+162.05)TV**NVTANAGDLTVGNGAEINATEGAATLTTSSGK |
| Site 2' | ATTGEANVTSATGTIGGTISGNTVNVTA**N(+162.05)AG**DLTVGNGAEINATEGAATLTTSSGK |
| Site 3' | ATTGEANVTSATGTIGGTISGNTVNVTANAGDLTVG**N(+162.05)GA**EINATEGAATLTTSSGK |
| Site 5' | GQVNLSAQDGSVAGSI**N(+162.05)AA**NVTLNTTGTLTTVK |
| Site 6' | GS**N(+162.05)IN**ATSGTLVINAK |
| Site 7' | GSNINATSGTLVI**N(+324.11)AK**DAELNGAALGNHTVVNATNANGSGSVIATTSSR |
| Site 8' | GSNINATSGTLVINAKDAEL**N(+162.05)GA**ALGNHTVVNATNANGSGSVIATTSSR |
| Site 9' | GSNINATSGTLVINAKDAELNGAALGNHTVVNAT**N(+162.05)AN**GSGSVIATTSSR |
